# The Influence of Obesity and Body Shape on Sagittal Plane Knee Kinematics and Kinetics during Obstacle Crossing

**DOI:** 10.64898/2026.09.24.753943

**Authors:** Chi-Whan Choi, Cara L. Lewis, Simone V. Gill

## Abstract

Altered walking mechanics in individuals with obesity can contribute to knee osteoarthritis. The gait deviations may become more pronounced during obstacle crossing. In women, body fat distribution may further influence knee load, especially when excess fat accumulates in the thighs and hips. However, relatively little is known about how regional fat distribution affects gait in women with obesity. This study investigated how obesity and, among women, different fat distributions (Apple: more abdominal fat; Pear: more lower-limb fat) influence knee biomechanics during walking with and without obstacle crossing. Participants were 15 controls without obesity (NB) and 27 with obesity (OB). Within female participants, 10 without obesity (fNB) were compared with 20 with obesity, stratified by waist-hip ratio (Apple:10, Pear:10). Speed-adjusted statistical parametric mapping applied a general linear model (NB vs. OB) and an analysis of covariance (fNB vs. Apple vs. Pear). OB exhibited a significantly greater late-stance knee extension moment than NB across all tasks, and this difference persisted among fNB, Apple, and Pear in obstacle tasks (*p*<0.05). OB walked with reduced knee flexion during the early-stance leading limb after crossing a medium-height obstacle (*p*=0.048) and a high-height obstacle (*p*=0.008) compared to NB. There were significant body-shape effects (*p*<0.05), and post-hoc comparisons confirmed that Pear had lower knee angles than fNB in both leading-limb conditions after crossing medium- and high-height obstacles (*p*=0.008 and *p*=0.001, respectively). These findings suggest that obstacle crossing helps illuminate how excess weight influences knee biomechanics, and how regional fat distribution modulates the degree of this alteration.

Keyword: Obesity; Body fat distribution; Osteoarthritis, Knee; Gait biomechanics; Obstacle crossing

## 1. Introduction

Obesity is associated with an increased risk and severity of knee osteoarthritis (KOA) (Li et al., 2022; Misra et al., 2019). Individuals with obesity have altered gait and make biomechanical adaptations (Del Porto et al., 2012; Kim et al., 2022), which may influence how forces are distributed across the knee joint. A study which explored kinematic differences at both preferred and fast gait speeds, showed that obesity was related to walking with an extended knee in early stance at the faster speed (Lerner et al., 2014a). Walking with a straighter leg may reflect a gait strategy to minimize vastus muscle force requirement (Vakula et al., 2022), as adults with obesity have been reported to have relatively weaker quadriceps function (Bollinger and Ransom, 2020).

Along with gait differences, obesity is correlated with sensorimotor dysfunction, involving both impaired motor planning and altered afferent input. Individuals with obesity show altered motor planning within the sensorimotor system due to managing biomechanical constraints imposed by increased adipose tissue (LoJacono et al., 2018). Altered afferent sensory signals may also contribute to poor sensorimotor processing given the reduced amplitude of stretch reflexes in muscles coupled with restricted range of motion (ROM) in individuals with obesity (Meng et al., 2017; Park et al., 2010). Although these deficits may remain latent during simple motor tasks, their impact would be expected to grow as tasks demand greater postural control and dynamic balance (Bourdon et al., 2026). Sensorimotor problems can exacerbate gait impairment when distracted or when confronted with tasks to complete (Mignardot et al., 2010), and altered gait that is linked with obesity is more evident when individuals with obesity encounter obstacles (Gill, 2019; Gill et al., 2016; Kim and Gill, 2020). For example, a previous study revealed that obstacle clearance performance is influenced by an individual’s body mass index (BMI) and the use of compensatory behaviors to regain stability (Lim et al., 2023).

Even though atypical gait induced by obesity has been comprehensively documented (Browning, 2012; Hills et al., 2002; Runhaar et al., 2011), only BMI was used to classify obesity. BMI, though widely used, often oversimplifies obesity because it does not account for regional fat distribution. When accounting for fat distribution, obesity is commonly categorized into two types: android obesity (apple-shaped, with fat concentrated in the abdominal region) and gynoid obesity (pear-shaped, with fat primarily in the lower extremities) (Ghezelbash et al., 2017). Body shape has often been overlooked in biomechanical studies, although body mass distribution can significantly influence stability and gait (Cieślińska-Świder et al., 2017; Maktouf et al., 2024). Women with obesity tend to accumulate a greater proportion of fat in the thighs, consistent with a gynoid pattern (Browning et al., 2006), and face a well-established risk for both the development and progression of KOA (Garcia et al., 2021; Misra et al., 2019). However, it remains unclear how regional fat distribution affects knee biomechanics that contribute to KOA risk in women with obesity. In dynamic activities such as obstacle crossing, whether fat distribution exerts a greater influence on knee biomechanics also remains to be determined.

Based on these gaps in the literature, we chose to investigate how obesity and differences in mass distribution affect knee biomechanics while walking at preferred and fast speeds on flat ground and when navigating obstacles in people with obesity compared to those without obesity. We hypothesized that (1) individuals with obesity would display altered knee kinematics and kinetics compared to controls with normal BMI across the walking tasks; and (2) among women, body shape would affect knee biomechanics, with pear-shaped women showing the most pronounced alterations, particularly during obstacle crossing.

## 2. Method

The methods are briefly described here with more details in the supplementary material.

### 2.1. Participants and protocol

A total of 42 participants were recruited: 15 adults (5 male; 10 female) with normal BMI and 27 adults (7 male; 20 female [apple shape (Apple):10, pear shape (Pear):10]) with obesity. The female participants with apple shape obesity were stratified according to waist-hip ratio (WHR) and a body shape index (ABSI) while the female participants with pear shape obesity were categorized with WHR and Hip index (HI) (Christakoudi et al., 2022). Since WHR indicates only the ratio and not the regional distribution, ABSI (abdominal) and HI (gluteal) were added to WHR to classify Apple and Pear more strictly. The classification cutoffs, eligibility criteria, and exclusion criteria are provided in the supplementary material. The study was approved by the Boston University Institutional Review Board (#7599) and conformed to the Declaration of Helsinki. Informed consent was obtained before testing began.

Participants were asked to wear tight fitting shorts and t-shirt. Since subcutaneous adipose tissue makes accurate measurement of kinematics particularly challenging in individuals with overweight and obesity, an obesity-specific marker set incorporating a sacral cluster was implemented (Lerner et al., 2014b) (Fig. S1, <u>supplemental video</u>). Six tasks were performed: preferred speed walking (PRF), fast speed walking (FF), stance of trailing limb (TL) before obstacle crossing medium height (OCMB-TL), stance of TL before obstacle crossing high height (OCHB-TL), stance of leading limb (LL) after obstacle crossing medium height (OCMA-LL), and stance of LL after obstacle crossing high height (OCHA-LL). The trailing and leading limbs were collected separately due to their distinct biomechanical roles during obstacle negotiation; the dominant leg was used for data analysis. Six participants reported the left leg as dominant (i.e., the preferred kicking leg), while the remaining participants reported right leg dominance. After participants became familiar with each task, participants walked across a 10m walk-way. Participants were allowed to rest as needed. Each task was repeated 10 times, and task order was randomized, except for the PRF, which was always completed first (Fig. 1).

**Fig. 1.**
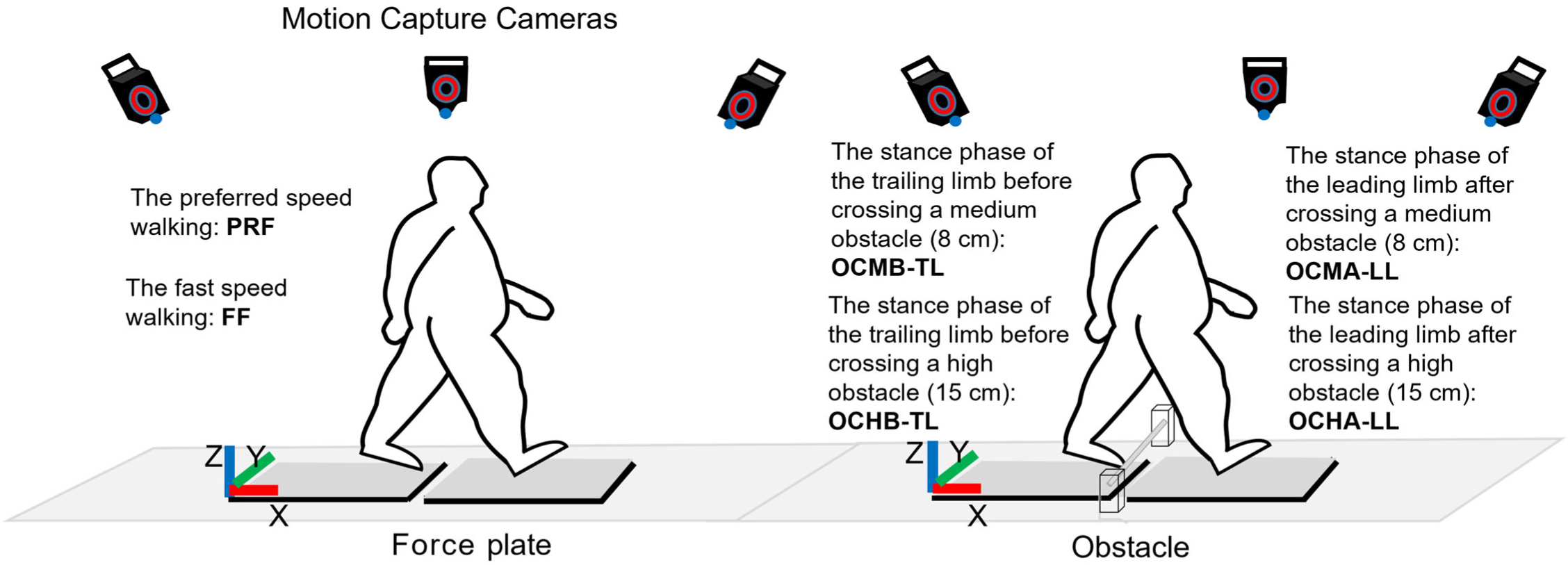
Experimental setup with motion capture system, and force plates while overground walking (Left), and walking with obstacle crossing (Right).

### 2.2. Data collection and processing

Three-dimensional trunk and lower extremity kinematic data were collected using a 10-camera motion-capture system (Vicon, Oxford, UK), consisting of 10 high-resolution cameras (sampling rate of 100 Hz) to track location of reflective markers placed on a participant. To obtain the kinetics of the walking, ground reaction force (GRF) data were recorded during walking with floor-embedded force platforms (Kistler force plate, type 9286AA, Kistler, Switzerland). The kinematic and kinetic data were synchronized using Nexus software (Oxford Metrics, Oxford, UK). All data were then imported into Visual3D (HAS Motion, Kingston, Ontario, Canada). A participant-specific, 8-segment hybrid model, including trunk, pelvis, bilateral thighs, shanks, and feet was created in Visual3D. Marker and GRF data were filtered using a low-pass Butterworth filter with cut-off frequencies of 6 Hz to remove noise and retain relevant signals. Each clean foot strike and each clean toe-off were identified from the GRF data (threshold:15 N). We identified the foot strike and toe-off events for the leg of interest, which mark the beginning and end of each stance cycle. The body segment parameters (mass, COM, and radius of gyration ratios) (de Leva, 1996) and GRF were used for the inverse-dynamics calculation of the internal knee moment. Moments were normalized to body mass (Nm/kg). At least four trials per task per participant (range: 4–10 trials, maximum 10 trials) were selected for data analysis, excluding trials with inaccurate foot placement on the force plate. Each walking speed for each trial was calculated using standard Visual3D commands, excluding strides during initial acceleration and final deceleration.

Walking speed was not controlled, as enforcing a fixed speed could disrupt natural obstacle avoidance. Instead, walkway length and obstacle location were fixed across trials within each task to promote consistent speed across trials. Time-continuous sagittal plane knee angle and knee moment were the primary outcome measures. The secondary outcome measures comprised the following discrete values. Peak knee angle (Peak Angle), knee flexion excursion from initial knee angle to peak knee angle (Knee Excursion), 1^st^ and 2^nd^ peak knee moment (Peak Moment), and knee moment range from 1^st^ peak flexion moment to 1^st^ peak extension moment (Moment Range) within each task (details in supplementary material).

### 2.3. Statistical analysis

Participant characteristics, anthropometry, and gait speed were compared using t-tests or Mann-Whitney U tests, depending on normality (Shapiro-Wilk test). Time-specific group differences in the kinematic and kinetic time-series were assessed with statistical parametric mapping (SPM). For the first aim, NB and OB were compared using a general linear model (GLM) with walking speed as a covariate, alongside an uncorrected GLM for reference. For the second aim, differences among fNB, Apple, and Pear were evaluated using an analysis of covariance (ANCOVA) with walking speed as a covariate, alongside an uncorrected analysis of variance (ANOVA) for reference. When significant differences were detected, post hoc t-tests with Benjamini-Hochberg correction (false discovery rate, FDR) were performed (Savage et al., 2021). All SPM analyses used the open-source SPM1D Python package (www.spm1d.org). For secondary analysis, random effects linear mixed models (LMMs) in R software (version 4.4.0) were implemented. To ensure consistent inference and account for potential heteroscedasticity and within-subject dependence, fixed effects were tested using cluster-robust standard errors (CR2), with clustering at the subject level (Pustejovs.ky and Tipton, 2018). Model performance was evaluated using Conditional *R^2^* (variance explained by fixed and random effects) and Marginal *R^2^*(variance explained by fixed effects alone).

## 3. Results

### 3.1. Participant characteristics

The general characteristics of the participants, group differences (NB vs. OB) in all participants, and body-shape differences (fNB vs. Apple vs. Pear) among female participants are presented in Table 1.

**Table 1.**
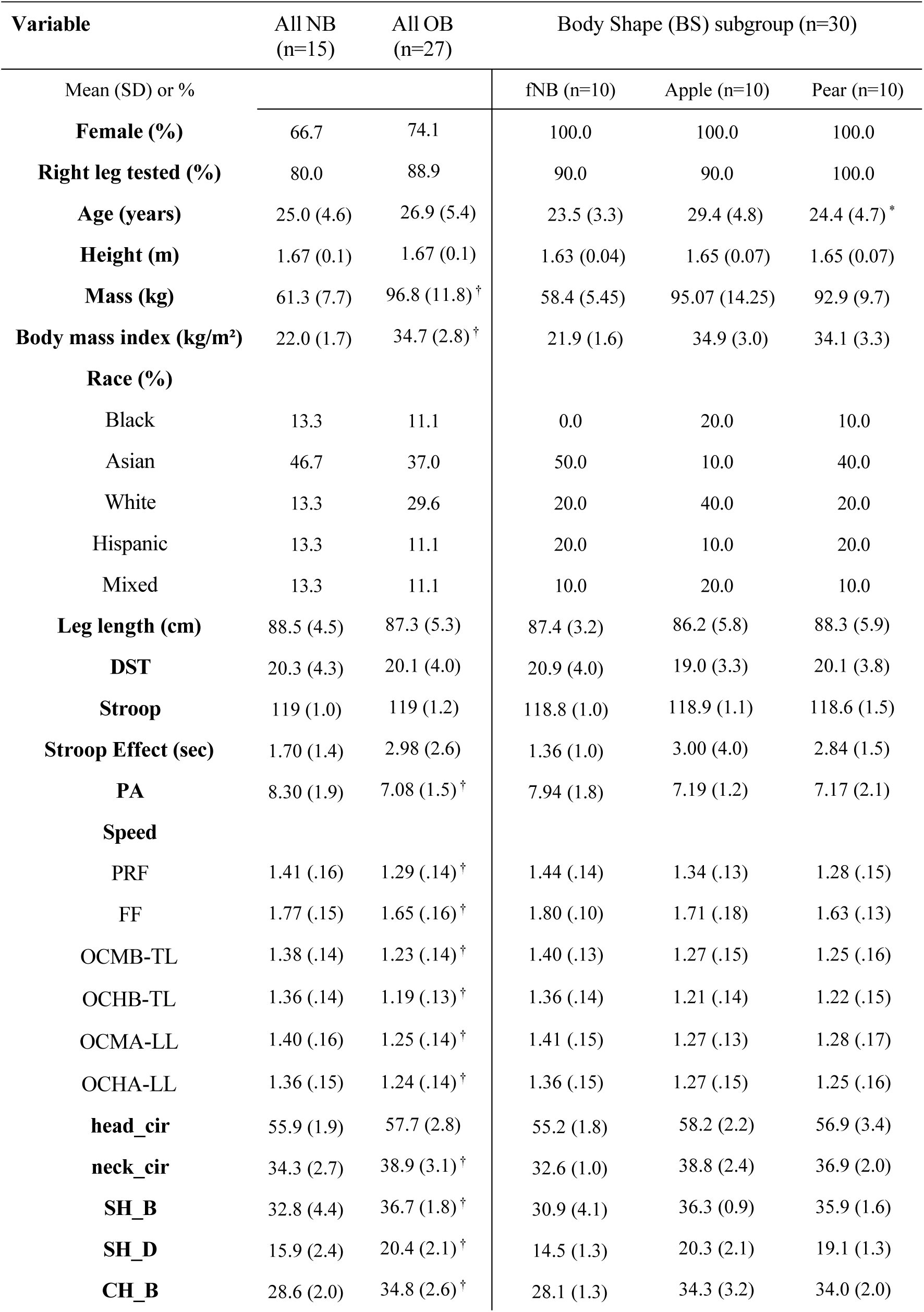
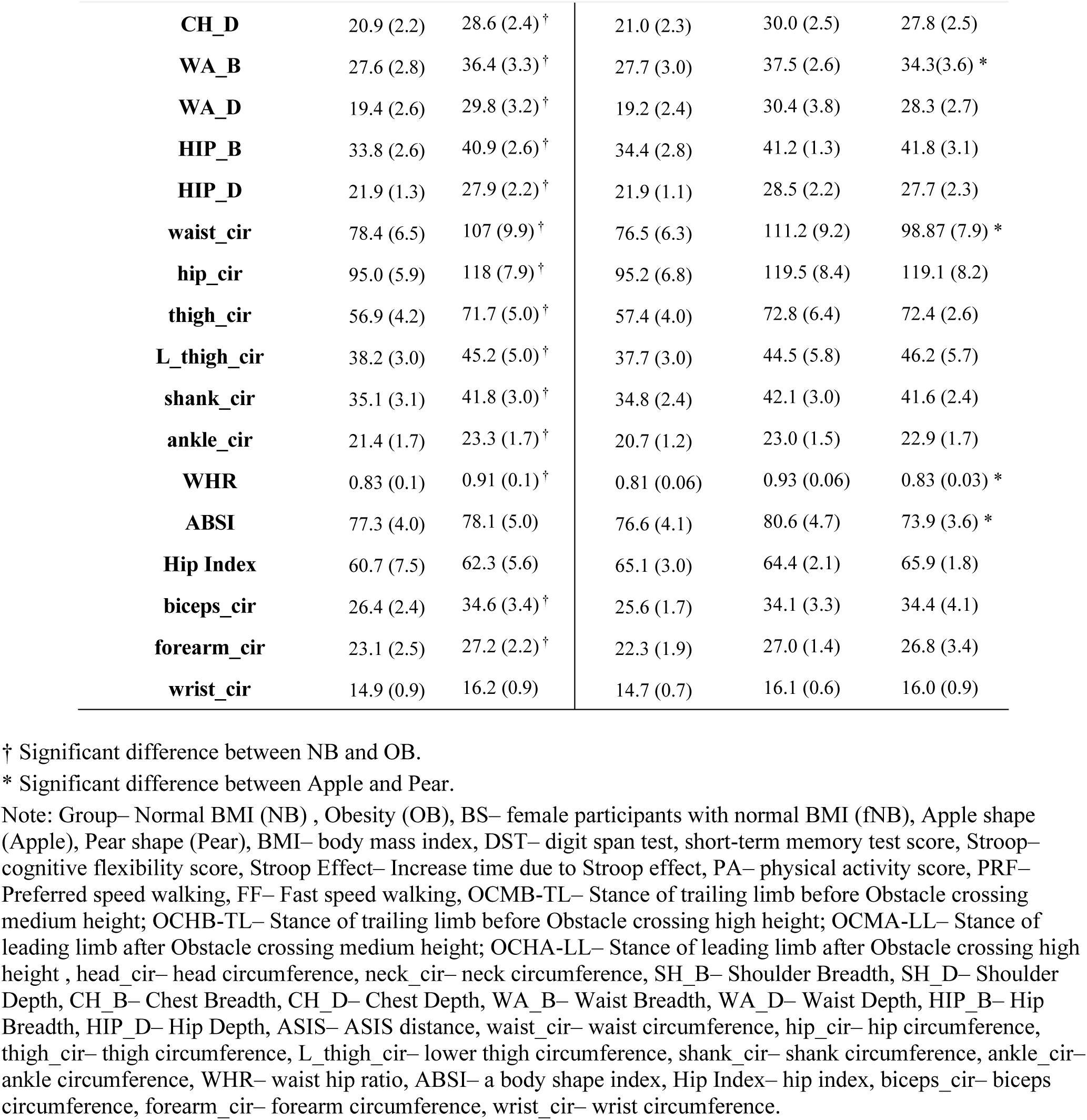
Participant characteristics (N = 42) for each group and each subgroup.

### 3.2. Group differences with time-continuous analysis, SPM GLM

In all tasks, OB exhibited significantly less knee flexion (all, *p* < 0.05) than NB during the initial contact, loading response or mid-stance regions of the stance phase (0–100%) before speed adjustments. After speed adjustment, significant group differences remained only in the OCMA-LL (10–14%; *p* = 0.048, Cohen’s *d (d)* = 1.194) and OCHA-LL (4–30%; *p* = 0.008, *d* = 1.271) (Fig. 2).

**Fig. 2.**
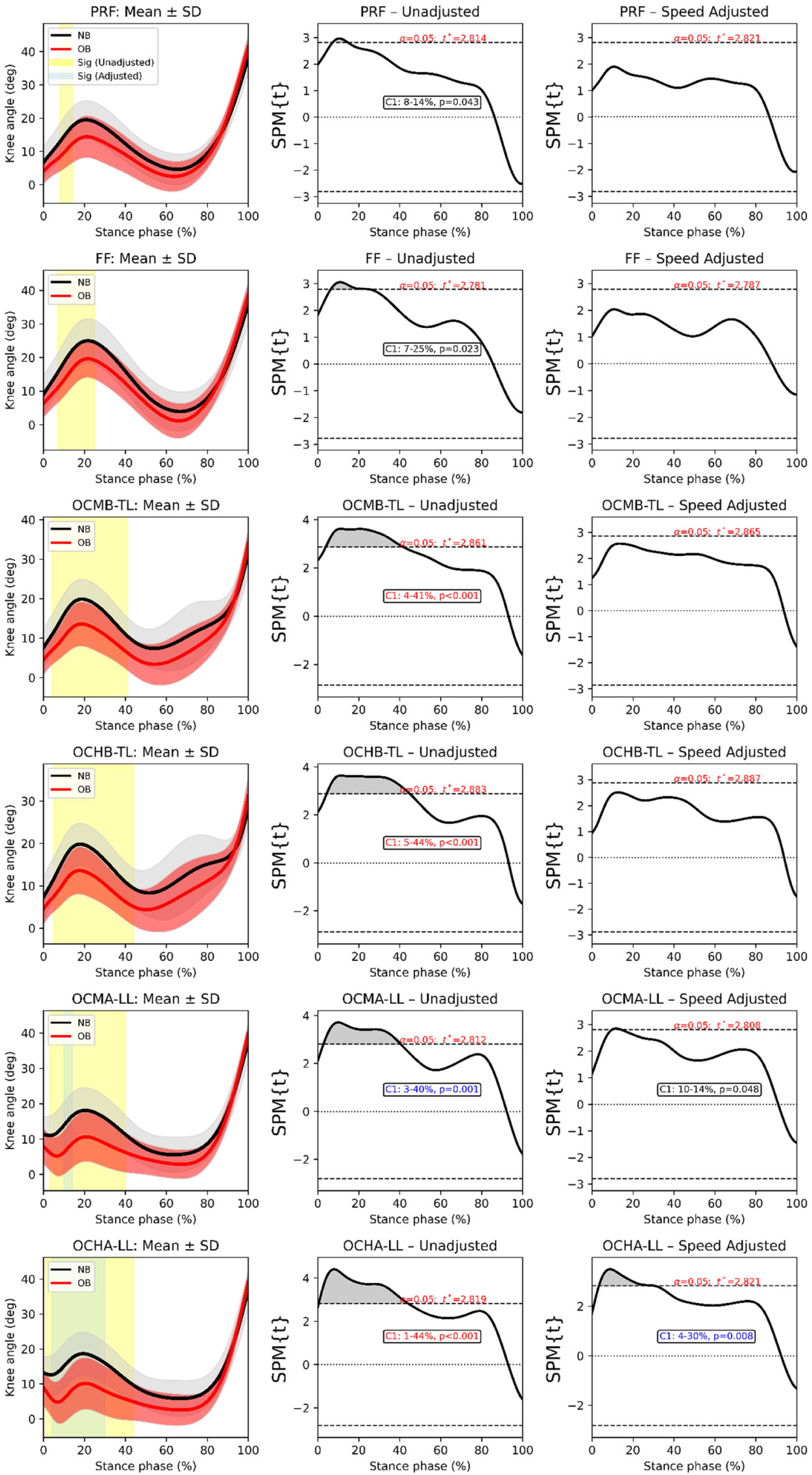
Average knee joint angle gait biomechanics showing standard deviation (SD) for Normal BMI (black) and Obese (red) groups within each task. Both a general linear model (GLM), with and without walking speed as covariates were applied using SPM. Also shown are regions where the statistical parametric map test statistic exceeded the critical threshold using i) a general linear model (yellow shading) without gait speed as covariates and ii) a general linear model (blue shading) with gait speed as covariates. The scalar output statistic of SPM, denoted SPM{t} for GLM was computed independently at each time point. SPM{t} was directly related to the magnitude of the difference between groups. Note: PRF– Preferred speed walking; FF– Fast speed walking; OCMB-TL– Stance of trailing limb before Obstacle crossing medium height; OCHB-TL– Stance of trailing limb before Obstacle crossing high height; OCMA-LL– Stance of leading limb after Obstacle crossing medium height; OCHA-LL– Stance of leading limb after Obstacle crossing high height; C1– First cluster; C2– Second cluster.

OB exhibited a significantly greater internal knee extension moment than NB during late stance across all tasks, and this difference persisted after speed adjustment (all, *p* < 0.05). An additional early-stance region showed a lower moment in OB in all tasks except for PRF, but these clusters were largely abolished by speed adjustment, remaining significant only in FF (11–16%, *p* = 0.030, *d* = 1.429) (Fig. 3).

**Fig. 3.**
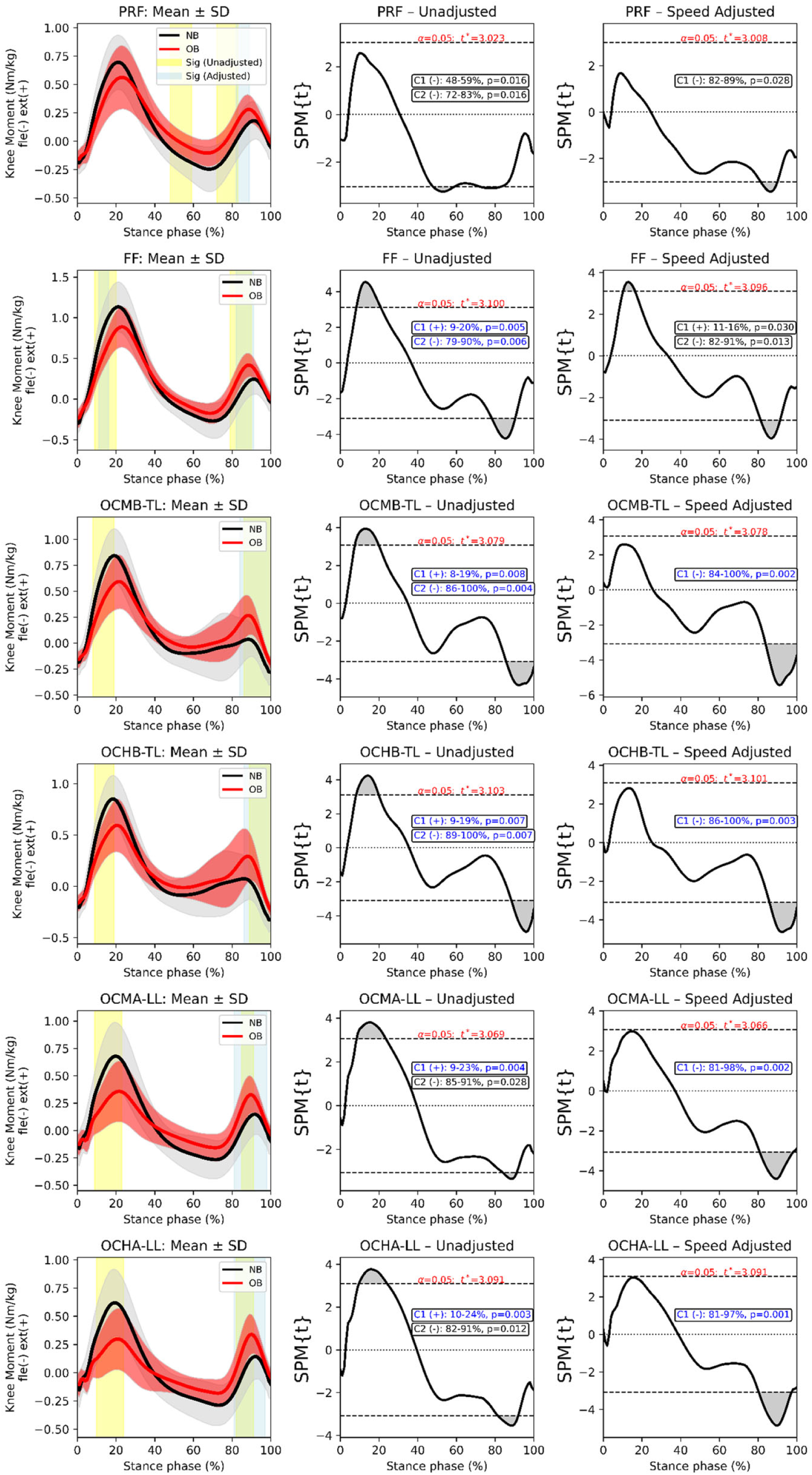
Average knee joint moment gait biomechanics showing standard deviation (SD) for Normal BMI (black) and Obese (red) groups within each task. Both a general linear model (GLM), with and without walking speed as covariates were applied using SPM. Also shown are regions where the statistical parametric map test statistic exceeded the critical threshold using i) a general linear model (yellow shading) without gait speed as covariates and ii) a general linear model (blue shading) with gait speed as covariates. The scalar output statistic of SPM, denoted SPM{t} for GLM was computed independently at each time point. SPM{t} was directly related to the magnitude of the difference between groups. Note: PRF– Preferred speed walking; FF– Fast speed walking; OCMB-TL– Stance of trailing limb before Obstacle crossing medium height; OCHB-TL– Stance of trailing limb before Obstacle crossing high height; OCMA-LL– Stance of leading limb after Obstacle crossing medium height; OCHA-LL– Stance of leading limb after Obstacle crossing high height; C1– First cluster; C2– Second cluster; (+)– NB > OB; (-)– NB < OB; Black– Significance with p <0.05; Blue– Significance with p <0.01; Red– Significance with p <0.001. Knee moment (Nm/kg) was normalized to body weight.

### 3.3. Body shape group differences with SPM ANOVA and ANCOVA

In all tasks, initial-to-mid stance knee flexion differed significantly among fNB, Apple, and Pear (all, *p* < 0.01) before speed adjustments. After speed adjustment, both OCMA-LL and OCHA-LL showed a significant early-stance group effect (OCMA-LL: 6–19%, *p* = 0.031; OCHA-LL: 1–32%, *p* = 0.003) (Fig. 4). Post-hoc comparisons showed that the group difference was driven by the fNB versus Pear comparison (OCMA-LL, *p* = 0.008; OCHA-LL, *p* = 0.001), which was the only comparison to remain significant after FDR correction (Table S1).

**Fig. 4.**
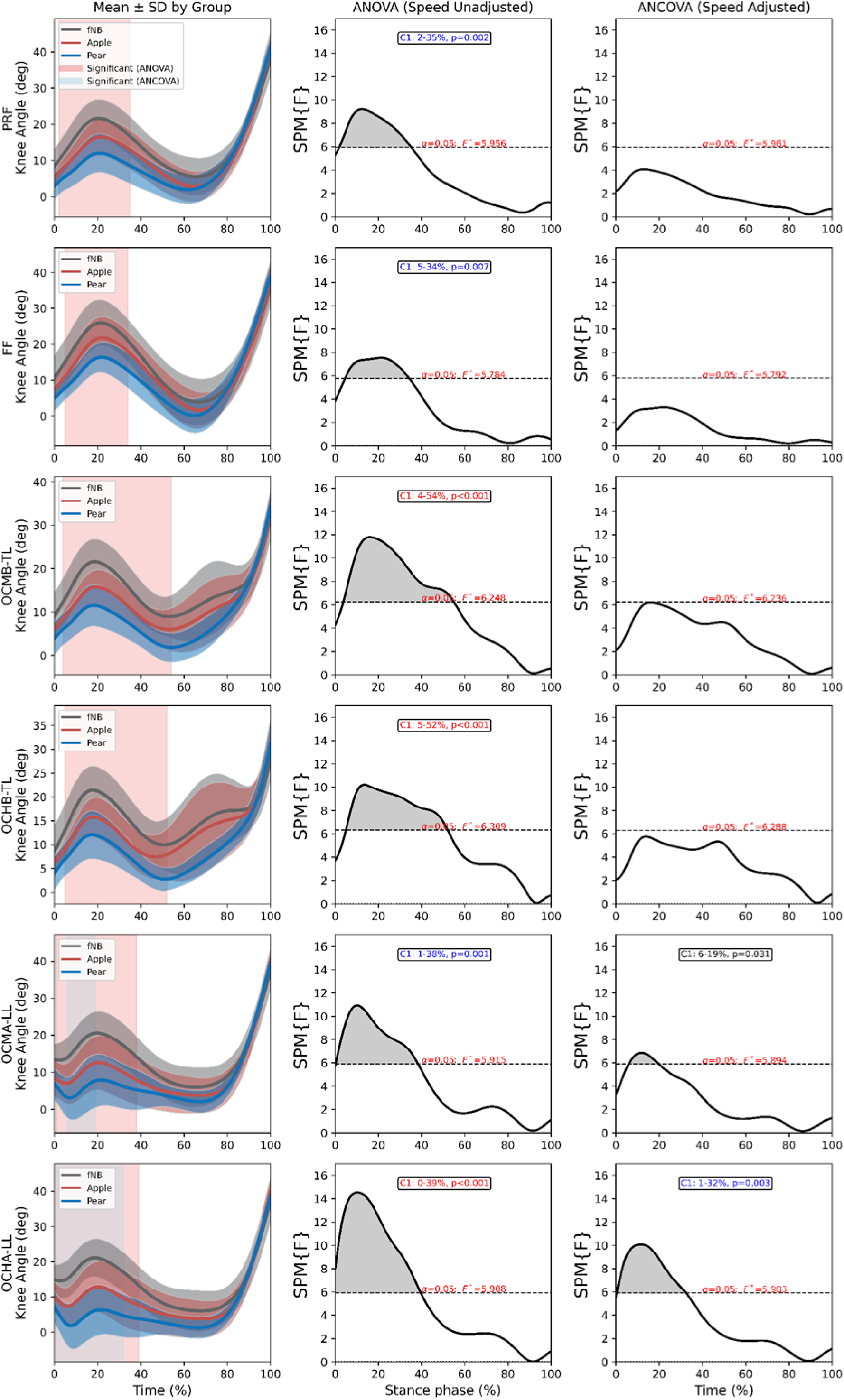
Average knee joint angle gait biomechanics showing standard deviation (SD) for female participants with normal BMI (fNB, black), apple shape (Apple, red), and pear shape (Pear, blue) groups within each task. Both ANOVA and ANCOVA with walking speed as covariates were applied using SPM. Also shown are regions where the statistical parametric map test statistic exceeded the critical threshold using i) ANOVA (pink shading) and ii) ANCOVA (blue shading) with gait speed as covariates. The scalar output statistic of SPM, denoted SPM{F} for ANOVA and ANCOVA was computed independently at each time point. SPM{F} was directly related to the magnitude of the difference between groups. Note: PRF– Preferred speed walking; FF– Fast speed walking; OCMB-TL– Stance of trailing limb before Obstacle crossing medium height; OCHB-TL– Stance of trailing limb before Obstacle crossing high height; OCMA-LL– Stance of leading limb after Obstacle crossing medium height; OCHA-LL– Stance of leading limb after Obstacle crossing high height; C1– First cluster; C2– Second cluster; Black – Significance with p <0.05; Blue – Significance with p <0.01; Red – Significance with p <0.001.

There were significant internal knee extension moment differences among fNB, Apple, and Pear in all tasks (all, *p* < 0.05) except for PRF. After speed adjustment, there were significant differences in all obstacle tasks during late stance (all, *p* < 0.05), with an additional early-stance difference in the OCHA-LL task (12–21%, *p* = 0.012) (Fig. 5). Post-hoc comparisons indicated that the knee moment differences were driven by the fNB versus Apple, and fNB versus Pear comparisons, which remained significant after FDR correction in all obstacle tasks (Table S2).

**Fig. 5.**
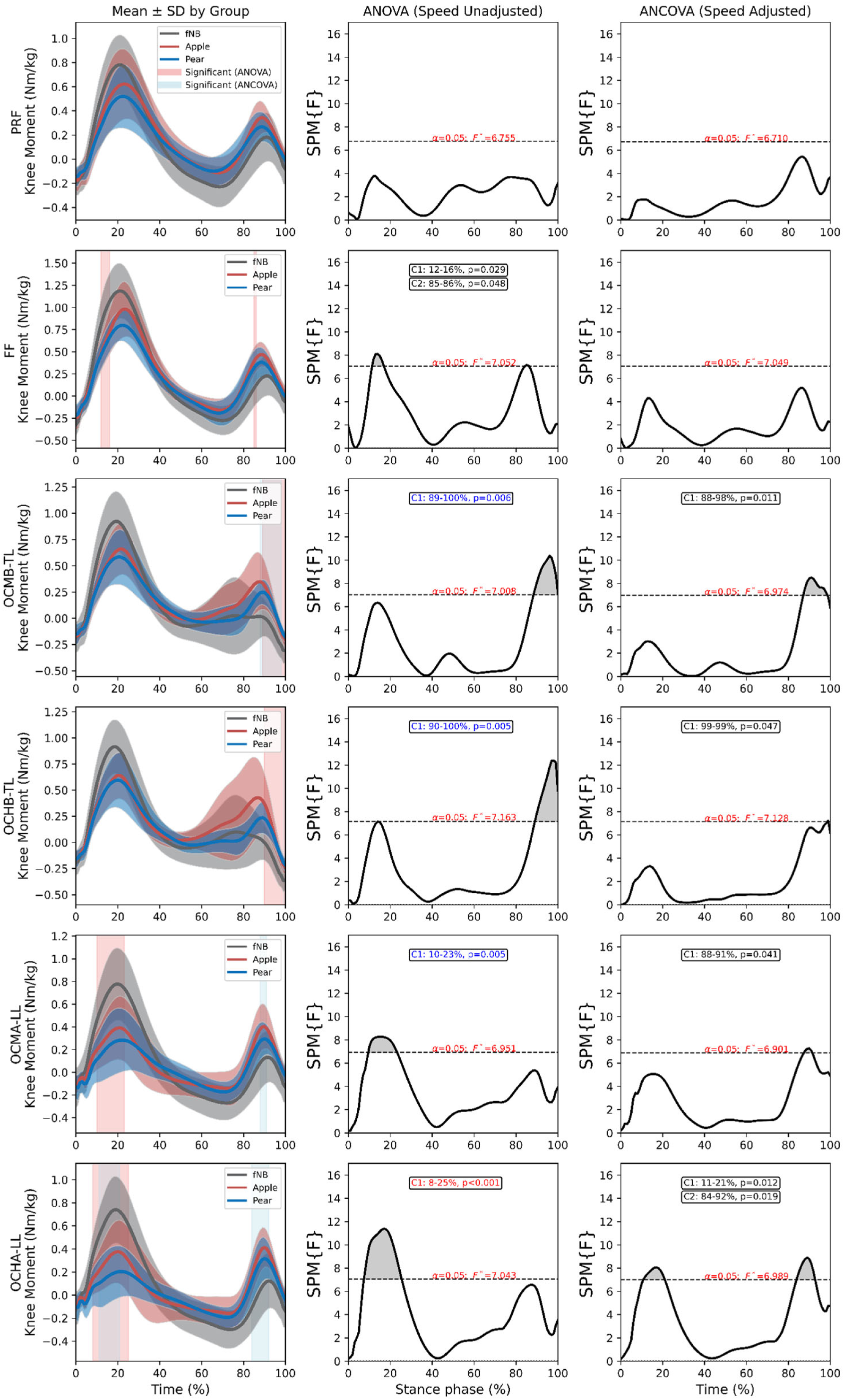
Average knee joint moment gait biomechanics showing standard deviation (SD) for female participants with normal BMI (fNB, black), apple shape (Apple, red), and pear shape (Pear, blue) groups within each task. Both ANOVA and ANCOVA with walking speed as covariates were applied using SPM. Also shown are regions where the statistical parametric map test statistic exceeded the critical threshold using i) ANOVA (pink shading) and ii) ANCOVA (blue shading) with gait speed as covariates. The scalar output statistic of SPM, denoted SPM{F} for ANOVA and ANCOVA was computed independently at each time point. SPM{F} was directly related to the magnitude of the difference between groups. Note: PRF– Preferred speed walking; FF– Fast speed walking; OCMB-TL– Stance of trailing limb before Obstacle crossing medium height; OCHB-TL– Stance of trailing limb before Obstacle crossing high height; OCMA-LL– Stance of leading limb after Obstacle crossing medium height; OCHA-LL– Stance of leading limb after Obstacle crossing high height. Knee moment (Nm/kg) was normalized to body weight; C1– First cluster; C2– Second cluster; Black – Significance with p <0.05; Blue – Significance with p <0.01; Red – Significance with p <0.001. Knee moment (Nm/kg) was normalized to body weight.

### 3.4. Discrete analysis with LMMs

LMMs revealed that OB walked with smaller Peak Angle (all tasks except PRF) and Knee Excursion (OCHB-TL, OCMA-LL, and OCHA-LL), a lower 1st Peak Moment and Moment Range (all tasks except PRF), and a higher 2nd Peak Moment (all tasks) than NB (Fig. 6). Full fixed-effect estimates, P values, and 95% confidence intervals (*CIs*) are provided in Table S3.

**Fig. 6.**
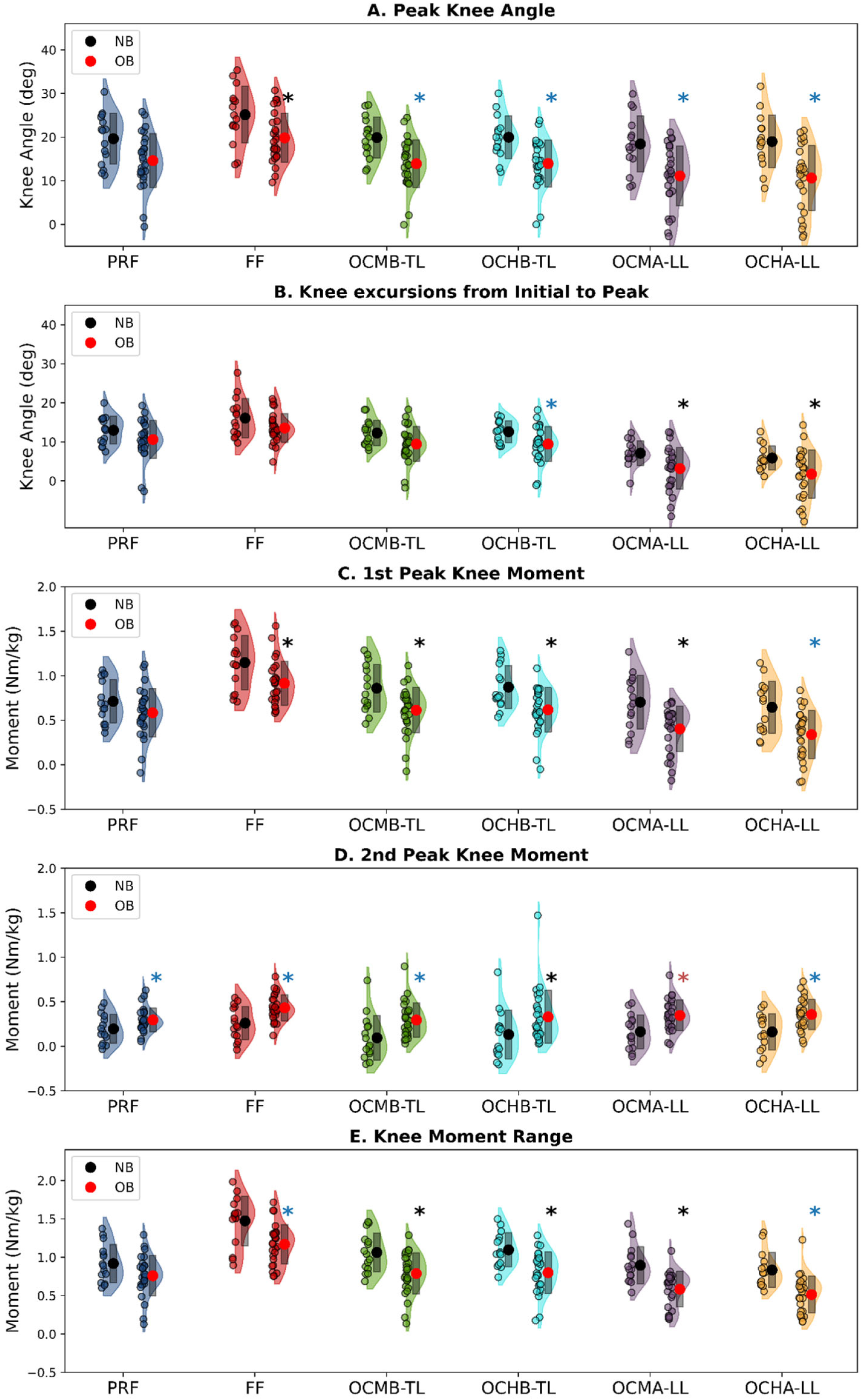
Rain cloud plots depicting mean distributions (shaded waveforms) for Normal BMI (NB, black), Obesity (OB, red). Different colors used to distinguish each task: PRF (dark blue), FF (dark red), OCMB-TL (green), OCHB-TL (cyan), OCMA-LL (purple), and OCHA-LL (orange). Note: PRF– Preferred speed walking; FF– Fast speed walking; OCMB-TL– Stance of trailing limb before Obstacle crossing medium height; OCHB-TL– Stance of trailing limb before Obstacle crossing high height; OCMA-LL– Stance of leading limb after Obstacle crossing medium height; OCHA-LL– Stance of leading limb after Obstacle crossing high height. Small circles represent data points from each individual participant, large circles and bars represent the data mean and standard deviation (SD) for each group. * – Significance with p <0.05; * – Significance with p <0.01; * – Significance with p <0.001. Knee moment (Nm/kg) was normalized to body weight.

Relative to fNB, Pear walked with lower Peak Angle and Knee Excursion (all tasks), lower 1^st^ Peak Moment and Moment Range (FF, OCMB-TL, OCMA-LL, and OCHA-LL), and greater 2^nd^ Peak Moment (FF, OCMA-LL, and OCHA-LL). Apple showed the same directional pattern but in fewer tasks: lower Peak Angle and Knee Excursion (OCHA-LL), lower 1^st^ Peak Moment (OCMA-LL, and OCHA-LL) and Moment Range (OCHA-LL), and greater 2^nd^ Peak Moment (FF, OCMB-TL, OCMA-LL, and OCHA-LL) (Fig. 7). Full fixed-effect estimates, *p* values, and 95% *CIs* are provided in Table S4.

**Fig. 7.**
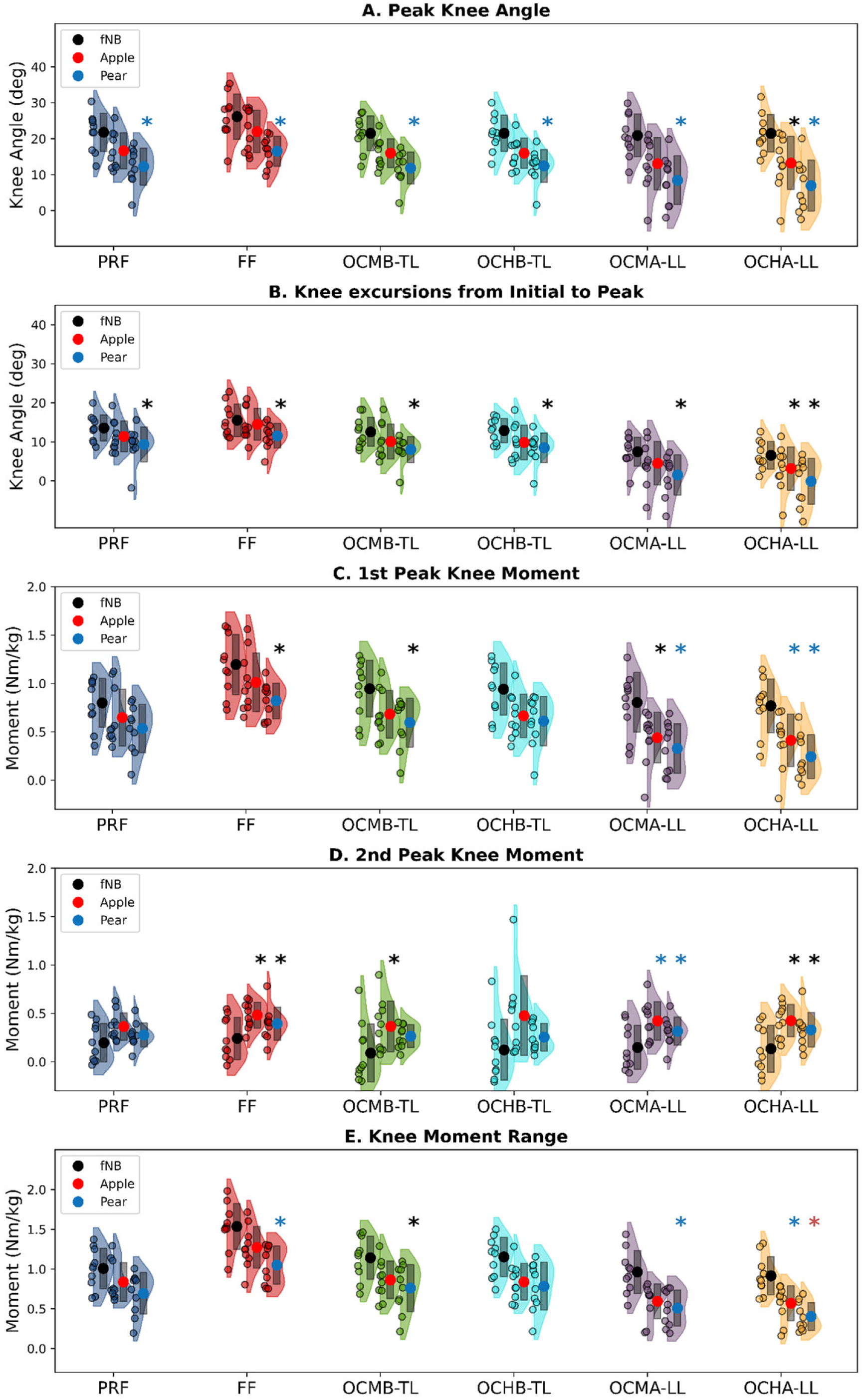
Rain cloud plots depicting mean distributions (shaded waveforms) for female participants with normal BMI (fNB, black), apple shape (Apple, red), pear shape (Pear, blue). Different colors used to distinguish each task: PRF (dark blue), FF (dark red), OCMB-TL (green), OCHB-TL (cyan), OCMA-LL (purple), and OCHA-LL (orange). Note: PRF– Preferred speed walking; FF– Fast speed walking; OCMB-TL– Stance of trailing limb before Obstacle crossing medium height; OCHB-TL– Stance of trailing limb before Obstacle crossing high height; OCMA-LL– Stance of leading limb after Obstacle crossing medium height; OCHA-LL– Stance of leading limb after Obstacle crossing high height. Small circles represent data points from each individual participant, large circles and bars represent the data mean and standard deviation (SD) for each group. * – Significance with p <0.05; * – Significance with p <0.01; * – Significance with p <0.001. Knee moment (Nm/kg) was normalized to body weight.

## 4. Discussion

This study examined sagittal knee biomechanics in participants with obesity (OB) compared to those with normal BMI (NB), as well as between female participants with different body shapes across level walking and obstacle crossing. In early stance, OB exhibited smaller knee flexion angles during the leading-limb stance after obstacle crossing and lower knee extension moments during fast walking compared to NB. OB demonstrated greater knee extension moments than NB in late stance across all tasks. Female participants with a pear shape (Pear) walked with reduced early-stance knee flexion angles compared to female participants with normal BMI (fNB) after obstacle crossing. Both Pear and those with an apple shape (Apple) showed greater knee extension moments in late stance than fNB in obstacle tasks. To our knowledge, this is the first study to show that obstacle crossing can clarify how obesity alters knee joint biomechanics and how regional fat distribution modulates the degree of this alteration. Because the early- and late-stance moments emerge from different muscular sources (quadriceps and plantarflexors, respectively), each phase is interpreted separately below.

The most robust finding of this study was the greater knee extension moment in OB than in NB during late stance (terminal-stance to pre-swing), which persisted across both Apple and Pear, before and after speed adjustment, in all walking tasks, and in both time-continuous (SPM) and discrete (LMMs) analyses. Given that plantarflexors during push-off contribute to the second peak of the sagittal knee moment (Sasaki and Neptune, 2010; Simonsen et al., 1997) and that propelling a heavier body amplifies this plantarflexor demand (Adouni et al., 2024a; Maktouf et al., 2020), the late-stance peak moment could partially reflect increased soleus activation. The uniarticular soleus, although it does not cross the knee, induces a net knee extension acceleration during stance through dynamic coupling of its plantarflexion force with GRF (Lenhart et al., 2014; Stewart et al., 2007), so the greater late-stance moment in OB may indicate greater soleus activation during push-off. Increased soleus activation can reduce tibiofemoral joint contact force; for example, previous research indicated that biofeedback retraining increasing soleus or decreasing gastrocnemius activation lowered late-stance knee contact force in both healthy adults (Hellman et al., 2025; Uhlrich et al., 2022) and individuals with KOA (Joyce et al., 2026). Together, it seems plausible that the increased knee extension moments observed in our study reflect greater soleus activation, potentially representing a compensatory strategy to modulate knee loading during late stance.

The reduced knee angle during early stance may be interpreted in a parallel manner. Because small changes in walking kinematics such as knee flexion angle can substantially alter the magnitude of knee joint forces, modifying movement and muscle coordination toward strategies that contribute less to knee loading may be an effective way to reduce knee joint contact forces during walking (Nagano et al., 2015; Sasaki and Neptune, 2010). The reduced knee angle could help modulate knee contact forces by maintaining the knee joint in straighter positions that aligned the vertical and anterior-posterior GRFs more closely with the center of the knee joint (Browning and Kram, 2007). Indeed, OB showed statistically lower peak knee flexion (Peak Angle) compared to NB across all tasks except for PRF. Walking with a straighter leg was accompanied with smaller 1^st^ internal knee extension moments (1^st^ Peak Moment), which may reflect less quadriceps activation (Vakula et al., 2022). Notably, walking speed had no significant effect on knee flexion excursion (Knee Excursion) during weight acceptance after crossing medium obstacles, and before and after crossing high obstacles based on the LMMs, indicating that the reduction of knee excursion reflects an adaptation to excess mass rather than to slower gait. This task-dependence may suggest that more demanding walking tasks accentuate obesity-related knee adaptations.

The biomechanical impact of different mass distributions on knee angles may be understood in terms of where excess mass is carried and its resulting inertial demands on the knee. The possible mechanism could be that individuals with obesity typically have greater thigh mass, thigh moment of inertia, shank mass, and shank moment of inertia compared to individuals with a normal BMI (Daniell et al., 2014; Merrill et al., 2019). Because our body-shape comparison was restricted to women, thigh-fat accumulation is particularly relevant here: women with obesity showed a greater ratio of thigh mass to total body mass compared to males with obesity measured by dual-energy X-ray absorptiometry (DEXA) (Browning et al., 2006). Within body-shape groups, this greater thigh-mass ratio is expected to be most pronounced in the pear shape, adding proximal-limb mass and increasing the limb’s inertia (Goossens et al., 2021). Given that a heavier, higher-inertia limb increases the load that must be eccentrically absorbed during weight acceptance, maintaining a straighter knee may lower the eccentric extensor demand on the quadriceps. These inertial properties may contribute to such compensatory strategies (reduced early-stance knee angle and knee excursion) to limit the eccentric demand and joint loading when landing the limb, particularly during leading-limb stance after obstacle crossing (Fig. S2, <u>supplemental video</u>). Consistent with this, only the fNB versus Pear contrast remained significant after FDR correction (Table S1).

Although obesity-related alterations in knee joint kinematics and kinetics may reduce overall knee load, they can at the same time redistribute and concentrate mechanical stress within the joint (Anan et al., 2024). In late stance, greater soleus activation has been shown to reduce overall tibiofemoral contact force (Hellman et al., 2025), but it can simultaneously elevate the external knee adduction moment, potentially exacerbating medial compartment loading. This suggests that compensatory muscle activation patterns intended to offload specific joints may inadvertently increase stress in other ways. In early stance, when knee flexion excursion is reduced, joint loads are distributed over a smaller area where the knee is maximally loaded, potentially increasing stress on knee structures at specific sites (Favre et al., 2014). A previous study simulated how knee motion affects not only knee muscle power absorption related to eccentric muscle activity, but also the knee adduction moment (Nagano et al., 2015). In line with this simulation, a reduced knee flexion excursion has been associated with a greater knee adduction moment impulse in a large cross-sectional sample (Kang et al., 2026).

Obesity affects the contact location mainly in the medial compartment (Adouni et al., 2024b; Li et al., 2022), which may partially explain the high prevalence of medial knee OA in people with obesity (Wei et al., 2019). The reduced early-stance knee flexion, combined with the reduced sagittal-plane moment range (Moment Range), reflects a “stiff-knee” pattern, in which a smaller range between the peak flexion and extension moments indicates increased dynamic knee stiffness. Such increased stiffness has been linked to altered knee contact forces and to the high propensity for post-traumatic osteoarthritis in individuals with anterior cruciate ligament (ACL) reconstruction (Garcia et al., 2023). Accordingly, the late-stance and early-stance adaptations may lead to greater medial compartment loading, heightening KOA vulnerability.

There are several limitations in this study. First, the presence of subcutaneous adipose tissue can lead to errors in joint kinematics. However, we followed a strict procedure based on prior work (Lerner et al., 2014b) to address these challenges by implementing an obesity-specific marker set. Second, we did not measure muscle activations or estimate joint contact forces. Future work using electromyography and musculoskeletal modeling is required to test whether a soleus-driven mechanism underlies the greater late-stance knee extension moment in obesity. Third, a limitation of this study is the unequal recruitment of male and female participants, with a higher number of women compared to men. To mitigate the effect, we included sex as a covariate in the LMMs. Lastly, internal knee moments relied on regression-based segment parameters derived from lean populations (de Leva, 1996); however, as stance-phase moments are dominated by GRF, their effect on our outcomes is likely limited. Nonetheless, DEXA (Browning, 2012) or an obesity-specific regression model (d’ Angelis et al., 2026) for estimating segment inertial parameters would improve accuracy.

In summary, increased mass and different body shapes were associated with significant alterations in knee joint kinematics and kinetics during various gait tasks. These differences were especially pronounced in obstacle-based activities, even after adjusting for walking speed, reflecting intrinsic movement strategies influenced by environmental contexts. This may indicate the alterations are not solely due to increased body mass, but rather due to muscle coordination to avoid knee loading (Runhaar et al., 2011; Vakula et al., 2022). Additionally, fat distribution, in particular in the lower limbs, was strongly associated with altered knee kinematics. This is not to underestimate the substantial influence of abdominal fat accumulation on gait function (Maktouf et al., 2024). Rather, the impact of thigh fat, given its location proximal to the knee joint, could make it a promising therapeutic target for walking (Messier et al., 2014). Further research is required to determine an optimal knee intervention that can prevent further damage to knee joint structures affected by obesity across body shapes and various real-world settings.

## CRediT authorship contribution statement

Chi-Whan Choi: Writing – original draft, Writing – review & editing, Visualization, Validation, Investigation, Formal analysis, Data curation, Funding acquisition, Methodology, Conceptualization. Cara L. Lewis: Writing – review & editing, Validation, Resources, Supervision, Methodology, Conceptualization. Simone V. Gill: Writing – review & editing, Validation, Supervision, Project administration, Methodology, Conceptualization.

## Acknowledgements

The authors acknowledge the contribution of all of those who participated in this study. This work was supported by the Dudley Allen Sargent Research Fund from Boston University (Choi, PI). This work was presented as a poster at APTA Combined Sections Meeting (CSM), 2026.

## Acknowledgment of Financial Support

Sargent College Research Grant, Boston University

## Reference

Adouni, M., Alkhatib, F., Hajji, R., Faisal, T.R., 2024a. Effects of overweight and obesity on lower limb walking characteristics from joint kinematics to muscle activations. Gait Posture 113, 337–344. 10.1016/j.gaitpost.2024.06.024

Adouni, M., Aydelik, H., Faisal, T.R., Hajji, R., 2024b. The effect of body weight on the knee joint biomechanics based on subject-specific finite element-musculoskeletal approach. Sci. Rep. 14, 13777. 10.1038/s41598-024-63745-x

Anan, M., Tokuda, K., Tanimoto, K., Sawada, T., 2024. The relationship between knee flexion excursion and mechanical stress during gait in medial knee osteoarthritis. Clin. Biomech. 112, 106180. 10.1016/j.clinbiomech.2024.106180

Bollinger, L.M., Ransom, A.L., 2020. The Association of Obesity With Quadriceps Activation During Sit-to-Stand. Phys. Ther. 100, 2134–2143. 10.30701/ijc.1535

Bourdon, A., Damm, L., Gasnier, A., Guyot, P., Cock, V.C. De, Hammouma, S., Coestier, B., Attalin, V., Janaqi, S., Bardy, B.G., 2026. Obesity is linked to impaired sensorimotor synchronization during walking but not tapping 1–10.

Browning, R.C., 2012. Locomotion Mechanics in Obese Adults and Children. Curr. Obes. Rep. 1, 152–159. 10.1007/s13679-012-0021-z

Browning, R.C., Baker, E.A., Herron, J.A., Kram, R., 2006. Effects of obesity and sex on the energetic cost and preferred speed of walking. J. Appl. Physiol. 100, 390–398. 10.1152/japplphysiol.00767.2005

Christakoudi, S., Riboli, E., Evangelou, E., Tsilidis, K.K., 2022. Associations of body shape index (ABSI) and hip index with liver, metabolic, and inflammatory biomarkers in the UK Biobank cohort. Sci. Rep. 12, 1–14. 10.1038/s41598-022-12284-4

Cieślińska-Świder, J., Furmanek, M.P., Błaszczyk, J.W., 2017. The influence of adipose tissue location on postural control. J. Biomech. 60, 162–169. 10.1016/j.jbiomech.2017.06.027

d’ Angelis, O., Choi, C.W., Sureshkumar, H., Merone, M., Gill, S. V., Song, S., 2026. Estimating Body Segment Properties for Adults Across Normal and Obese Body Types : A Data-Driven Geometric Framework. arXiv Prepr. 10.64898/2026.07.03.736346

Daniell, N., Olds, T., Tomkinson, G., 2014. Volumetric differences in body shape among adults with differing body mass index values: An analysis using three-dimensional body scans. Am. J. Hum. Biol. 26, 156–163. 10.1002/ajhb.22490

de Leva, P., 1996. Adjustments to zatsiorsky-seluyanov’s segment inertia parameters. J. Biomech. 29, 1223–1230. 10.1016/0021-9290(95)00178-6

Del Porto, H.C., Pechak, C.M., Smith, D.R., Reed-Jones, R.J., 2012. Biomechanical Effects of Obesity on Balance. Int. J. Exerc. Sci. 5, 301–320.

Favre, J., Erhart-Hledik, J.C., Andriacchi, T.P., 2014. Age-related differences in sagittal-plane knee function at heel-strike of walking are increased in osteoarthritic patients. Osteoarthr. Cartil. 22, 464–471. 10.1016/j.joca.2013.12.014

Garcia, S.A., Johnson, A.K., Brown, S.R., Washabaugh, E.P., Krishnan, C., Palmieri-Smith, R.M., 2023. Dynamic knee stiffness during walking is increased in individuals with anterior cruciate ligament reconstruction. J. Biomech. 146, 111400. 10.1016/j.jbiomech.2022.111400

Garcia, S.A., Vakula, M.N., Holmes, S.C., Pamukoff, D.N., 2021. The influence of body mass index and sex on frontal and sagittal plane knee mechanics during walking in young adults. Gait Posture 83, 217–222. 10.1016/j.gaitpost.2020.10.010

Ghezelbash, F. and O.S.M.I.S.B.A.S.-S.R.A.M., Shirazi-Adl, A., Plamondon, A., Arjmand, N., Parnianpour, M., 2017. Obesity and Obesity Shape Markedly Influence Spine Biomechanics: A Subject-Specific Risk Assessment Model. Ann. Biomed. Eng. 45, 2373–2382. 10.1007/s10439-017-1868-7

Gill, S. V., 2019. Effects of obesity class on flat ground walking and obstacle negotiation. J. Musculoskelet. Neuronal Interact. 19, 448–454.

Gill, S. V., Walsh, M.K., Pratt, J.A., Toosizadeh, N., Najafi, B., Travison, T.G., 2016. Changes in spatiotemporal gait patterns during flat ground walking and obstacle crossing 1 year after bariatric surgery. Surg. Obes. Relat. Dis. 12, 1080–1085. 10.1016/j.soard.2016.03.029

Goossens, G.H., Jocken, J.W.E., Blaak, E.E., 2021. Sexual dimorphism in cardiometabolic health: the role of adipose tissue, muscle and liver. Nat. Rev. Endocrinol. 17, 47–66. 10.1038/s41574-020-00431-8

Hellman, E., Arokoski, J., Franz, J., Skipper, M., Kristian, R., Stenroth, L., 2025. Acute effects of soleus EMG biofeedback training on tibiofemoral joint contact forces in young healthy adults. J. Biomech. 184, 112646. 10.1016/j.jbiomech.2025.112646

Hills, A.P., Hennig, E.M., Byrne, N.M., Steele, J.R., 2002. The biomechanics of adiposity - Structural and functional limitations of obesity and implications for movement. Obes. Rev. 3, 35–43. 10.1046/j.1467-789X.2002.00054.x

Joyce, M.R., Muccini, J., Randoing, B., Delp, S.L., Uhlrich, S.D., 2026. Retraining Gastrocnemius Muscle Coordination Reduces Late-Stance Knee Contact Force in Individuals With Knee Osteoarthritis. IEEE Trans. Neural Syst. Rehabil. Eng. 34, 1417–1425. 10.1109/TNSRE.2026.3669842

Kang, K.S., Lee, N.K., Lee, K.M., Park, M.S., Chang, C.B., 2026. Increased knee flexion excursion during early stance is associated with reduced knee adduction moment in asymptomatic varus lower extremities: a cross-sectional study of 937 limbs. J. Biomech. 205, 113467. 10.1016/j.jbiomech.2026.113467

Kim, D., Gill, S. V., 2020. Changes in center of pressure velocities during obstacle crossing one year after bariatric surgery. Gait Posture 76, 377–381. 10.1016/j.gaitpost.2019.12.020

Kim, D., Lewis, C.L., Silverman, A.K., Gill, S. V., 2022. Changes in dynamic balance control in adults with obesity across walking speeds. J. Biomech. 144, 111308. 10.1016/j.jbiomech.2022.111308

Lenhart, R.L., Francis, C.A., Lenz, A.L., Thelen, D.G., 2014. Empirical evaluation of gastrocnemius and soleus function during walking. J. Biomech. 47, 2969–2974. 10.1016/j.jbiomech.2014.07.007

Lerner, Z.F., Board, W.J., Browning, R.C., 2014a. Effects of obesity on lower extremity muscle function during walking at two speeds. Gait Posture 39, 978–984. 10.1016/j.gaitpost.2013.12.020

Lerner, Z.F., Board, W.J., Browning, R.C., 2014b. Effects of an obesity-specific marker set on estimated muscle and joint forces in walking. Med. Sci. Sports Exerc. 46, 1261–1267. 10.1249/MSS.0000000000000218

Li, J., Tsai, T., Clancy, M.M., Lewis, C.L., Felson, D.T., Li, G., 2022. Cartilage contact characteristics of the knee during gait in individuals with obesity. J. Orthop. Res. 1–8. 10.1002/jor.25288

Lim, S., Luo, Y., Lee-Confer, J., D’Souza, C., 2023. Obstacle clearance performance in individuals with high body mass index. Appl. Ergon. 106, 103879. 10.1016/j.apergo.2022.103879

LoJacono, C.T., MacPherson, R.P., Kuznetsov, N.A., Raisbeck, L.D., Ross, S.E., Rhea, C.K., 2018. Obstacle crossing in a virtual environment transfers to a real environment. J. Mot. Learn. Dev. 6, 234–249. 10.1123/jmld.2017-0019

Maktouf, W., Durand, S., Boyas, S., Pouliquen, C., Beaune, B., 2020. Interactions among obesity and age-related effects on the gait pattern and muscle activity across the ankle joint. Exp. Gerontol. 140, 111054. 10.1016/j.exger.2020.111054

Maktouf, W., Ferhi, H., Boyas, S., Beaune, B., Chortane, S.G., Portero, P., Durand, S., 2024. The influence of obesity and fat distribution on ankle muscle coactivation during gait. PLoS One 19, 1–16. 10.1371/journal.pone.0294692

Meng, H., O’Connor, D.P., Lee, B.C., Layne, C.S., Gorniak, S.L., 2017. Alterations in over-ground walking patterns in obese and overweight adults. Gait Posture 53, 145–150. 10.1016/j.gaitpost.2017.01.019

Merrill, Z., Perera, S., Chambers, A., Cham, R., 2019. Age and body mass index associations with body segment parameters. J. Biomech. 88, 38–47. 10.1016/j.jbiomech.2019.03.016

Messier, S.P., Beavers, D.P., Loeser, R.F., Carr, J.J., Khajanchi, S., Legault, C., Nicklas, B.J., Hunter, D.J., Devita, P., 2014. Knee joint loading in knee osteoarthritis: Influence of abdominal and thigh fat. Med. Sci. Sports Exerc. 46, 1677–1683. 10.1249/MSS.0000000000000293

Mignardot, J.B., Olivier, I., Promayon, E., Nougier, V., 2010. Obesity impact on the attentional cost for controlling posture. PLoS One 5, 1–6. 10.1371/journal.pone.0014387

Misra, D., Fielding, R.A., Felson, D.T., Niu, J., Brown, C., Nevitt, M., Lewis, C.E., Torner, J., Neogi, T., 2019. Risk of Knee Osteoarthritis With Obesity, Sarcopenic Obesity, and Sarcopenia. Arthritis Rheumatol. 71, 232–237. 10.1002/art.40692

Nagano, H., Tatsumi, I., Sarashina, E., Sparrow, W.A., Begg, R.K., 2015. Modelling knee flexion effects on joint power absorption and adduction moment. Knee 22, 490–493. 10.1016/j.knee.2015.06.016

Park, W., Ramachandran, J., Weisman, P., Jung, E.S., 2010. Obesity effect on male active joint range of motion. Ergonomics 53, 102–108. 10.1080/00140130903311617

Pustejovsky, J.E., Tipton, E., 2018. Small-Sample Methods for Cluster-Robust Variance Estimation and Hypothesis Testing in Fixed Effects Models. J. Bus. Econ. Stat. 36, 672–683. 10.1080/07350015.2016.1247004

Runhaar, J., Koes, B.W., Clockaerts, S., Bierma-Zeinstra, S.M.A., 2011. A systematic review on changed biomechanics of lower extremities in obese individuals: A possible role in development of osteoarthritis. Obes. Rev. 12, 1071–1082. 10.1111/j.1467-789X.2011.00916.x

Sasaki, K., Neptune, R.R., 2010. Individual muscle contributions to the axial knee joint contact force during normal walking. J. Biomech. 43, 2780–2784. 10.1016/j.jbiomech.2010.06.011

Savage, T.N., Saxby, D.J., Pizzolato, C., Diamond, L.E., Murphy, N.J., Hall, M., Spiers, L., Eyles, J., Killen, B.A., Suwarganda, E.K., Dickenson, E.J., Griffin, D., Fary, C., O’Donnell, J., Molnar, R., Randhawa, S., Reichenbach, S., Tran, P., Wrigley, T. V., Bennell, K.L., Hunter, D.J., Lloyd, D.G., 2021. Trunk, pelvis and lower limb walking biomechanics are similarly altered in those with femoroacetabular impingement syndrome regardless of cam morphology size. Gait Posture 83, 26–34. 10.1016/j.gaitpost.2020.10.002

Simonsen, E.B., Dyhre-Poulsen, P., Voigt, M., Aagaard, P., Fallenlin, N., 1997. Mechanisms contributing to different joint moments observed during human walking. Scand. J. Med. Sci. Sport. 7, 1–13. 10.1111/j.1600-0838.1997.tb00110.x

Stewart, C., Postans, N., Schwartz, M.H., Rozumalski, A., Roberts, A., 2007. An exploration of the function of the triceps surae during normal gait using functional electrical stimulation. Gait Posture 26, 482–488. 10.1016/j.gaitpost.2006.12.001

Uhlrich, S.D., Jackson, R.W., Seth, A., Kolesar, J.A., Delp, S.L., 2022. Muscle coordination retraining inspired by musculoskeletal simulations reduces knee contact force. Sci. Rep. 12, 1–13. 10.1038/s41598-022-13386-9

Vakula, M.N., Garcia, S.A., Holmes, S.C., Pamukoff, D.N., 2022. Association between quadriceps function, joint kinetics, and spatiotemporal gait parameters in young adults with and without obesity. Gait Posture 92, 421–427. 10.1016/j.gaitpost.2021.12.019

Wei, J., Gross, D., Lane, N.E., Lu, N., Wang, M., Zeng, C., Yang, T., Lei, G., Choi, H.K., Zhang, Y., 2019. Risk factor heterogeneity for medial and lateral compartment knee osteoarthritis : analysis of two prospective cohorts. Osteoarthr. Cart. 27, 603–610. 10.1016/j.joca.2018.12.013

