## supplementary material for "The Influence of Obesity and Body Shape on Sagittal Plane Knee Kinematics and Kinetics during Obstacle Crossing"

**Method**

*Body shape stratification and eligibility criteria*

The female participants with apple shape obesity were stratified according to waist-hip ratio (WHR) and a body shape index (ABSI), with cut-offs WHR ≥ 0.85 and ABSI ≥ 73. The female participants with pear shape obesity were categorized with WHR and Hip index (HI), with cut-offs WHR < 0.85 and HI ≥ 64 [1]. Eligibility criteria included being between 18 and 39 years of age; having a body mass index (BMI) within 18.5 ≤ BMI < 25 kg/m² or 30 ≤ BMI < 40 kg/m², corresponding to normal weight or class I–II obesity, respectively, based on the Centers for Disease Control and Prevention classification[2]; being able to walk for at least 30 minutes without an assistive device; and being able to speak and understand English. Exclusion criteria were having other serious health problems such as cardiovascular disease, metabolic disease, neurological disease, head trauma, musculoskeletal pain, visual impairment or hearing impairment that would make it difficult to participate in the study.

*Sample size*

The sample size was based on a power analysis using G*Power (effect size f= 0.50, α = .05, power = 0.80).

*Procedure and protocol*

After consenting, we asked a participant's their age and sex. Participants' height and weight were measured with a wall-mounted stadiometer, and a weighing scale. BMI was calculated with the formula weight (kg)/height (m)$^2$. Anthropometric measurements including head circumference, neck base circumference, shoulder breadth, shoulder depth, chest breadth, chest depth, waist depth, waist breadth, hip breadth, hip depth, hip circumference, thigh circumference, lower thigh circumference, shank circumference, ankle circumference, biceps circumference, forearm circumference, and wrist circumference were measured to investigate the fat distribution patterns using an anthropometer and a tape measure. Leg length from the anterior superior iliac spine to the medial malleolus using tape measure was measured. Participants completed the Baecke Physical Activity Questionnaire [3]. Next, participants were asked to perform the Digit Span Task to measure working memory and attention, then they performed the Stroop test to examine cognitive flexibility.

Since subcutaneous adipose tissue makes accurate measurement of kinematics particularly challenging in individuals with overweight and obesity [4–6], an obesity-specific marker set incorporating a sacral cluster was implemented (Fig.S1). Lerner et al. [4] investigated the effects of the cluster of four markers and showed more accurate and repeatable measurements of pelvic kinematics, especially for overweight and obese subjects where soft tissue artifacts can be more pronounced [4,5]. Total of 51 reflective markers were placed on bony landmarks on the participant's trunk and lower extremities. Markers were placed bilaterally on the posterior heel, three metatarsal heads (1st, 3rd, and 5th), medial and lateral malleoli, medial and lateral femoral epicondyles, greater trochanter, iliac crest, anterior superior iliac spine (ASIS), between ASIS and iliac crest, between iliac crest and sacrum, and acromion process. A single marker was placed on the jugular notch, 7th cervical vertebra, and right scapular inferior angle. Five rigid clusters of four markers were placed via neoprene wraps on the bilateral thighs and shanks, and the sacrum. After placing reflective markers, the subjects were asked to stand still while left and right ASIS were determined using the tip of the calibration wand. The wand's technical coordinate frame was then used to define the position of each ASIS with respect to the coordinate frame of the cluster, called as Calibrated Anatomical System Technique (CAST) (Fig. S1, supplemental video)[7]. During dynamic trials, the positions of the ASIS and hip joint center were reconstructed based on their relationship to the sacral cluster defined in the static calibration. By using this technique, we aimed to minimize the effects of soft tissue artifact. Several short calibrations trials were collected throughout data collection. Each of these trials did not exceed 15 seconds.

Total six tasks including preferred speed walking (PRF), fast speed walking (FF), stance of trailing limb (TL) before obstacle crossing medium height (OCMB-TL), stance of TL before obstacle crossing high height (OCHB-TL), stance of leading limb (LL) after obstacle crossing medium height (OCMA-LL), and stance of LL after obstacle crossing high height (OCHA-LL) were performed.

- Preferred speed walking (PRF): Participants walked over floor-embedded force platforms with a 10-camera motion-capture across a 10m walk-way.
- Fast speed walking (FF): Participants walked across a 10m walk-way at 125% preferred speed.
- Walking with obstacle crossing (OCMB-TL, OCHB-TL, OCMA-LL, and OCHA-LL): Participants walked over obstacles of different heights. The obstacle heights were 8cm, and 15cm, and were held by wooden blocks with insertion points of a collapsing wood bar.

*Secondary outcome measures*

The secondary outcome measures comprised the following discrete values. Discrete extraction of such values is a conventional and reproducible approach in gait biomechanics. All are established variables from prior knee gait-analysis literature [8–16].

- Peak knee angle (Peak Angle): The maximum sagittal-plane knee flexion angle during the stance phase of each task. Peak knee flexion angle is a standard descriptor of weight-acceptance kinematics and has been used extensively to characterize the reduced-flexion (stiff-knee) gait in individuals with obesity [8], knee osteoarthritis [9], and anterior cruciate ligament (ACL) reconstruction [10].

- Knee flexion excursion (Knee Excursion): The angular displacement from the knee angle at initial contact to the peak stance knee flexion angle (i.e., early-stance flexion excursion) [9]. Knee flexion excursion during weight acceptance is a well-established loading-related variable: it is reduced in individuals with obesity [12] and is positively associated with peak knee extension moment.
- 1st and 2nd peak knee moment (Peak Moment): The first (early-stance) and second (late-stance) peaks of the internal sagittal knee moment. This variable is a standard kinetic measure extracted in obesity[14], osteoarthritis[15] and ACL gait analyses[10].
- Knee moment range (Moment Range): The change in the internal knee moment between the 1st peak internal knee flexion moment and the 1st peak internal knee extension moment during early stance. This range reflects the change in sagittal knee moment during loading response and, when coupled with the corresponding angular excursion, underlies the widely used dynamic-joint-stiffness construct linked to knee osteoarthritis risk [16].

*Statistical analysis*

We used statistical parametric mapping (SPM) to identify time-specific group differences in kinematic and kinetic time-series data. SPM treats these one-dimensional trajectories as vector fields and applies random field theory to test whether observed variations between groups are statistically significant (i.e., unlikely to arise from chance). This approach minimizes biases associated with regional focus (e.g., peak-value analysis) and intercomponent covariance in continuous biomechanical data. The scalar output statistic of SPM, denoted SPM{t} for a general linear model (GLM) and SPM{F} for analysis of variance (ANOVA) and analysis of covariance (ANCOVA) was computed independently at each time point. SPM{t} and SPM{F} were directly related to the magnitude of the difference between groups. For secondary analysis, random effects linear mixed models (LMMs) in R software (version 4.4.0) were implemented. LMMs have the advantage of handling complex, multilevel data structures and analyzing repeated measures data. Thus, LMMs can account for within-subject correlations by incorporating random effects such as a random intercept for each subject. Group (NB vs OB) and potential confounding factors including age, sex, race, leg length and walking speed were entered into the model as fixed effects according to literature [9,17,18], and each participant was modeled as a random effect on the outcomes. Finally, we explored how Group (NB vs OB) and the confounding factors influenced the discrete outcome measures within each task. We also assessed how different body shapes (fNB vs Apple vs Pear) in female participants and the confounding factors except for sex influenced the same variables within each task.

**Reference**

1. Christakoudi S, Riboli E, Evangelou E, Tsilidis KK. Associations of body shape index (ABSI) and hip index with liver, metabolic, and inflammatory biomarkers in the UK Biobank cohort. *Sci Rep*. 2022;12(1):1-14. doi:10.1038/s41598-022-12284-4
2. Cornier MA, Despre JP, Davis N, et al. Assessing Adiposity. *Circulation*. 2011;124(18):1996-2019. doi:10.1161/CIR.0b013e318233bc6a
3. Tebar WR, Ritti-Dias RM, Fernandes RA, et al. Validity and reliability of the Baecke questionnaire against accelerometermeasured physical activity in community dwelling adults according to educational level. *PLoS One*. 2022;17(8 August):1-11. doi:10.1371/journal.pone.0270265
4. Lerner ZF, Board WJ, Browning RC. Effects of an obesity-specific marker set on estimated muscle and joint forces in walking. *Med Sci Sports Exerc*. 2014;46(6):1261-1267. doi:10.1249/MSS.0000000000000218
5. Borhani M, McGregor AH, Bull AMJ. An alternative technical marker set for the pelvis is more repeatable than the standard pelvic marker set. *Gait Posture*. 2013;38(4):1032-1037. doi:10.1016/j.gaitpost.2013.05.019
6. Horsak B, Pobatschnig B, Baca A, et al. Within-assessor reliability and minimal detectable change of gait kinematics in a young obese demographic. *Gait Posture*. 2017;54:112-118. doi:10.1016/j.gaitpost.2017.02.028
7. Aurelio C. Gait analysis methodology. *Hum Mov Sci*. 1984;3(1-2):27-50.
8. Lerner ZF, Board WJ, Browning RC. Effects of obesity on lower extremity muscle function during walking at two speeds. *Gait Posture*. 2014;39(3):978-984. doi:10.1016/j.gaitpost.2013.12.020

9. Favre J, Erhart-Hledik JC, Andriacchi TP. Age-related differences in sagittal-plane knee function at heel-strike of walking are increased in osteoarthritic patients. *Osteoarthr Cartil*. 2014;22(3):464-471. doi:10.1016/j.joca.2013.12.014

10. Krishnan C, Johnson AK, Palmieri-Smith RM. Mechanical Factors Contributing to Altered Knee Extension Moment during Gait after ACL Reconstruction: A Longitudinal Analysis. *Med Sci Sports Exerc*. 2022;54(12):2208-2215. doi:10.1249/MSS.0000000000003014

11. Adouni M, Alkhatib F, Hajji R, Faisal TR. Effects of overweight and obesity on lower limb walking characteristics from joint kinematics to muscle activations. *Gait Posture*. 2024;113(June):337-344. doi:10.1016/j.gaitpost.2024.06.024

12. Pamukoff DN, Lewek MD, Blackburn JT. Greater vertical loading rate in obese compared to normal weight young adults. *Clin Biomech*. 2016;33:61-65. doi:10.1016/j.clinbiomech.2016.02.007

13. Kang KS, Lee NK, Lee KM, Park MS, Chang CB. Increased knee flexion excursion during early stance is associated with reduced knee adduction moment in asymptomatic varus lower extremities: a cross-sectional study of 937 limbs. *J Biomech*. 2026;205(July):113467. doi:10.1016/j.jbiomech.2026.113467

14. Davis-Wilson HC, Johnston CD, Young E, et al. Effects of BMI on Walking Speed and Gait Biomechanics after Anterior Cruciate Ligament Reconstruction. *Med Sci Sports Exerc*. 2021;53(1):108-114. doi:10.1249/MSS.0000000000002460

15. Davis HC, Luc-Harkey BA, Seeley MK, Troy Blackburn J, Pietrosimone B. Sagittal plane walking biomechanics in individuals with knee osteoarthritis after quadriceps strengthening. *Osteoarthr Cartil*. 2019;27(5):771-780. doi:10.1016/j.joca.2018.12.026

16. Garcia SA, Johnson AK, Brown SR, Washabaugh EP, Krishnan C, Palmieri-Smith RM. Dynamic knee stiffness during walking is increased in individuals with anterior cruciate ligament reconstruction. *J Biomech*. 2023;146(November 2022):111400. doi:10.1016/j.jbiomech.2022.111400

17. Garcia SA, Vakula MN, Holmes SC, Pamukoff DN. The influence of body mass index and sex on frontal and sagittal plane knee mechanics during walking in young adults. *Gait Posture*. 2021;83(September 2020):217-222. doi:10.1016/j.gaitpost.2020.10.010

18. Hill CN, Reed W, Schmitt D, Arent SM, Sands LP, Queen RM. Factors contributing to racial differences in gait mechanics differ by sex. *Gait Posture*. 2022;95(February 2021):277-283. doi:10.1016/j.gaitpost.2021.02.024

**Supplemental Table S1.** Post-Hoc Pairwise Comparisons for OCMA-LL and OCHA-LL after ANCOVA.

| | Task | Comparison | Corrected *p*-value (FDR) | Cohen's *d* | 95% *CI* (Lower, Upper) |
|---|---|---|---|---|---|
| Knee Angle | OCMA-LL | fNB vs Apple | 0.074 | 0.941 | 0.156 – 2.011 |
| | | fNB vs Pear | **0.008**** | 1.574 | 0.779 – 2.962 |
| | | Apple vs Pear | 0.121 | 0.730 | -0.101 – 1.856 |
| | OCHA-LL | fNB vs Apple | 0.524 | 1.020 | 0.282 – 1.991 |
| | | fNB vs Pear | **0.001**** | 1.899 | 1.083 – 3.543 |
| | | Apple vs Pear | 0.061 | 0.894 | 0.044 – 2.249 |

Note: fNB (female participants with normal BMI); Apple shape (Apple); Pear shape (Pear); OCMA-LL– Stance of leading limb after Obstacle crossing medium height; OCHA-LL– Stance of leading limb after Obstacle crossing high height; FDR– False discovery rate control; *CI*– Confidence interval for Cohen's d; * – Significance with p <0.05; ** – Significance with p <0.01.

**Supplemental Table S2.** Post-Hoc Pairwise Comparisons for OCMB-TL, OCHM-TL, OCMA-LL, and OCHA-LL after ANCOVA.

| | Task | Comparison | Corrected *p*-value (FDR) | Cohen's *d* | 95% CI (Lower, Upper) |
|---|---|---|---|---|---|
| Knee Moment | OCMB-TL | fNB vs Apple | **0.005**** | -1.498 | -2.818 – -0.712 |
| | | fNB vs Pear | **0.005**** | -1.656 | -3.260 – -0.830 |
| | | Apple vs Pear | 0.648 | 0.209 | -0.734 – 1.185 |
| | OCHB-TL | fNB vs Apple | **0.025*** | -1.193 | -2.897 – -0.325 |
| | | fNB vs Pear | **0.011*** | -1.543 | -3.628 – -0.616 |
| | | Apple vs Pear | 0.518 | -0.295 | -1.250 – 0.635 |
| | OCMA-LL | fNB vs Apple | **0.010*** | -1.510 | -3.058 – -0.684 |
| | | fNB vs Pear | **0.040*** | -1.074 | -2.674 – -0.250 |
| | | Apple vs Pear | 0.160 | 0.658 | -0.225 – 1.680 |
| | OCHA-LL (1st cluster) | fNB vs Apple | **0.032*** | 1.131 | 0.276 – 2.742 |
| | | fNB vs Pear | **0.005**** | 1.675 | 0.765 – 3.758 |
| | | Apple vs Pear | 0.224 | 0.613 | -0.249 – 1.795 |
| | OCHA-LL (2nd cluster) | fNB vs Apple | **0.005**** | -1.727 | -3.509 – -0.873 |
| | | fNB vs Pear | **0.024*** | -1.255 | -2.731 – -0.451 |
| | | Apple vs Pear | 0.315 | 0.462 | -0.374 – 1.707 |

Note: fNB (female participants with normal BMI group); Apple shape (Apple); Pear shape (Pear); OCMA-LL– Stance of leading limb after Obstacle crossing medium height; OCHA-LL– Stance of leading limb after Obstacle crossing high height; FDR– False discovery rate control; CI– Confidence interval for Cohen's d; * – Significance with p <0.05; ** – Significance with p <0.01.

**Supplemental Table S3.** The effects of predictors on knee discrete outcomes in each task (PRF, FF, OCMB-TL, OCHB-TL, OCMA-LL and OCHA-LL) in all participants.

| | Source | *β* | *p* | *95%CI lower for β* | *95%CI upper for β* |
|---|---|---|---|---|---|
| **PRF** | (Intercept) | 9.358 | 0.661 | -36.54 | 55.25 |
| | **group** | -2.711 | 0.113 | -6.114 | 0.693 |
| | **age** | -0.015 | 0.945 | -0.471 | 0.441 |
| Peak Angle | **sex** | 1.076 | 0.669 | -4.126 | 6.278 |
| | **leg length** | -0.135 | 0.511 | -0.575 | 0.306 |
| | **race** | | | | |
| | Black | Ref. | Ref. | Ref. | |
| | Asian | 3.193 | 0.290 | -3.469 | 9.855 |
| | White | -2.777 | 0.366 | -9.484 | 3.931 |
| | Hispanic | 1.950 | 0.461 | -3.839 | 7.739 |
| | Mixed | 0.739 | 0.766 | -4.747 | 6.225 |
| | **speed** | 14.43 | **<.001*****| 10.25 | 18.62 |
| *Conditional $R^2$* | 0.956 | | | | |
| *Marginal $R^2$* | 0.330 | | | | |
| | (Intercept) | 16.51 | 0.273 | -15.11 | 48.14 |
| | **group** | -0.519 | 0.702 | -3.283 | 2.246 |
| | **age** | -0.149 | 0.455 | -0.563 | 0.265 |
| Knee Excursion | **sex** | -1.212 | 0.577 | -5.694 | 3.269 |
| | **leg length** | -0.153 | 0.345 | -0.496 | 0.190 |
| | **race** | | | | |
| | Black | Ref. | Ref. | Ref. | |
| | Asian | 0.165 | 0.937 | -4.644 | 4.974 |
| | White | -2.655 | 0.259 | -7.702 | 2.391 |
| | Hispanic | 0.799 | 0.766 | -5.155 | 6.753 |
| | Mixed | 0.366 | 0.861 | -4.241 | 4.973 |
| | **speed** | 10.39 | **0.003**** | 3.828 | 16.96 |
| *Conditional $R^2$* | 0.899 | | | | |
| *Marginal $R^2$* | 0.207 | | | | |
| | (Intercept) | 0.955 | 0.199 | -0.588 | 2.497 |
| | **group** | -0.015 | 0.814 | -0.142 | 0.113 |
| | **age** | -0.006 | 0.475 | -0.023 | 0.011 |
| 1st Peak Moment | **sex** | 0.004 | 0.970 | -0.209 | 0.216 |
| | **leg length** | -0.013 | 0.081 | -0.029 | 0.002 |
| | **race** | | | | |
| | Black | Ref. | Ref. | Ref. | |
| | Asian | 0.028 | 0.783 | -0.203 | 0.258 |
| | White | -0.152 | 0.175 | -0.388 | 0.084 |
| | Hispanic | 0.152 | 0.145 | -0.064 | 0.368 |
| | Mixed | 0.116 | 0.385 | -0.172 | 0.403 |
| | **speed** | 0.748 | **<.001*****| 0.467 | 1.030 |
| *Conditional $R^2$* | 0.919 | | | | |
| *Marginal $R^2$* | 0.382 | | | | |
| | (Intercept) | -0.749 | **0.039*** | -1.451 | -0.048 |
| | **group** | 0.125 | **0.023*** | 0.019 | 0.231 |
| | **age** | 0.009 | **0.027*** | 0.001 | 0.017 |
| 2nd Peak Moment | **sex** | 0.036 | 0.401 | -0.052 | 0.124 |
| | **leg length** | 0.002 | 0.551 | -0.005 | 0.010 |
| | **race** | | | | |
| | Black | Ref. | Ref. | Ref. | |
| | Asian | 0.013 | 0.776 | -0.095 | 0.122 |
| | White | 0.021 | 0.717 | -0.110 | 0.153 |

| | | | | | |
|---|---|---|---|---|---|
| | Hispanic | 0.051 | 0.402 | -0.082 | 0.185 |
| | Mixed | -0.056 | 0.383 | -0.193 | 0.082 |
| | **speed** | 0.356 | **<.001*****| 0.191 | 0.521 |
| *Conditional $R^2$* | 0.927 | | | | |
| *Marginal $R^2$* | 0.320 | | | | |
| Moment Range | (Intercept) | 0.698 | 0.326 | -0.802 | 2.198 |
| | **group** | -0.004 | 0.939 | -0.120 | 0.112 |
| | **age** | -0.006 | 0.448 | -0.021 | 0.010 |
| | **sex** | -0.028 | 0.760 | -0.218 | 0.162 |
| | **leg length** | -0.013 | 0.072 | -0.028 | 0.001 |
| | **race** | | | | |
| | Black | Ref. | Ref. | Ref. | |
| | Asian | -0.006 | 0.946 | -0.206 | 0.194 |
| | White | -0.164 | 0.088 | -0.359 | 0.031 |
| | Hispanic | 0.107 | 0.302 | -0.115 | 0.330 |
| | Mixed | 0.029 | 0.797 | -0.219 | 0.278 |
| | **speed** | 1.113 | **<.001*****| 0.847 | 1.379 |
| *Conditional $R^2$* | 0.929 | | | | |
| *Marginal $R^2$* | 0.267 | | | | |
| **FF** Peak Angle | (Intercept) | 18.53 | 0.357 | -24.06 | 61.12 |
| | **group** | -4.872 | **0.028*** | -9.156 | -0.587 |
| | **age** | 0.223 | 0.303 | -0.223 | 0.670 |
| | **sex** | -0.143 | 0.948 | -4.701 | 4.415 |
| | **leg length** | -0.112 | 0.532 | -0.498 | 0.275 |
| | **race** | | | | |
| | Black | Ref. | Ref. | Ref. | |
| | Asian | 5.990 | 0.113 | -1.892 | 13.87 |
| | White | 1.408 | 0.707 | -6.974 | 9.790 |
| | Hispanic | 4.465 | 0.191 | -2.742 | 11.67 |
| | Mixed | 7.445 | 0.062 | -0.452 | 15.34 |
| | **speed** | 3.635 | **0.025*** | 0.504 | 6.766 |
| *Conditional $R^2$* | 0.943 | | | | |
| *Marginal $R^2$* | 0.286 | | | | |
| Knee Excursion | (Intercept) | 25.23 | 0.046 | 0.498 | 49.95 |
| | **group** | -2.682 | 0.079 | -5.697 | 0.332 |
| | **age** | 0.173 | 0.328 | -0.192 | 0.538 |
| | **sex** | -2.314 | 0.204 | -6.004 | 1.376 |
| | **leg length** | -0.176 | 0.114 | -0.402 | 0.051 |
| | **race** | | | | |
| | Black | Ref. | Ref. | Ref. | |
| | Asian | 2.774 | 0.267 | -2.730 | 8.277 |
| | White | 1.730 | 0.475 | -3.616 | 7.076 |
| | Hispanic | 3.615 | 0.163 | -1.812 | 9.043 |
| | Mixed | 6.392 | **0.023*** | 1.119 | 11.66 |
| | **speed** | 0.421 | 0.727 | -2.050 | 2.891 |
| *Conditional $R^2$* | 0.872 | | | | |
| *Marginal $R^2$* | 0.225 | | | | |
| 1st Peak Moment | (Intercept) | 0.832 | 0.229 | -0.608 | 2.271 |
| | **group** | -0.182 | **0.042*** | -0.358 | -0.007 |
| | **age** | 0.006 | 0.432 | -0.009 | 0.021 |
| | **sex** | -0.007 | 0.943 | -0.195 | 0.181 |
| | **leg length** | -0.009 | 0.161 | -0.023 | 0.004 |
| | **race** | | | | |
| | Black | Ref. | Ref. | Ref. | |
| | Asian | 0.265 | 0.014 | 0.074 | 0.455 |
| | White | 0.099 | 0.250 | -0.086 | 0.285 |
| | Hispanic | 0.338 | **0.012*** | 0.097 | 0.58 |
| | Mixed | 0.386 | **0.032*** | 0.041 | 0.731 |
| | **speed** | 0.434 | **0.002**** | 0.184 | 0.684 |

| | | | | | |
|---|---|---|---|---|---|
| *Conditional $R^2$* | 0.896 | | | | |
| *Marginal $R^2$* | 0.393 | | | | |
| 2nd Peak Moment | (Intercept) | -1.209 | 0.022* | -2.205 | -0.212 |
| | **group** | 0.215 | 0.002** | 0.091 | 0.340 |
| | **age** | 0.008 | 0.143 | -0.003 | 0.020 |
| | **sex** | -0.021 | 0.713 | -0.142 | 0.099 |
| | **leg length** | 0.007 | 0.157 | -0.003 | 0.017 |
| | **race** | | | | |
| | Black | Ref. | Ref. | Ref. | |
| | Asian | 0.019 | 0.766 | -0.129 | 0.167 |
| | White | -0.021 | 0.793 | -0.201 | 0.158 |
| | Hispanic | 0.105 | 0.328 | -0.127 | 0.337 |
| | Mixed | -0.072 | 0.339 | -0.234 | 0.090 |
| | **speed** | 0.375 | <.001*** | 0.221 | 0.529 |
| *Conditional $R^2$* | 0.882 | | | | |
| *Marginal $R^2$* | 0.328 | | | | |
| Moment Range | (Intercept) | 0.487 | 0.473 | -0.959 | 1.934 |
| | **group** | -0.216 | 0.009** | -0.371 | -0.06 |
| | **age** | 0.008 | 0.273 | -0.007 | 0.022 |
| | **sex** | -0.021 | 0.811 | -0.205 | 0.163 |
| | **leg length** | -0.008 | 0.220 | -0.023 | 0.006 |
| | **race** | | | | |
| | Black | Ref. | Ref. | Ref. | |
| | Asian | 0.282 | 0.031 | 0.036 | 0.527 |
| | White | 0.066 | 0.527 | -0.165 | 0.297 |
| | Hispanic | 0.339 | 0.020 | 0.069 | 0.609 |
| | Mixed | 0.334 | 0.017 | 0.076 | 0.591 |
| | **speed** | 0.740 | <.001*** | 0.496 | 0.984 |
| *Conditional $R^2$* | 0.860 | | | | |
| *Marginal $R^2$* | 0.519 | | | | |
| **OCMB-TL** | (Intercept) | 12.16 | 0.489 | -25.38 | 49.70 |
| | **group** | -4.730 | 0.007** | -8.063 | -1.397 |
| | **age** | 0.057 | 0.777 | -0.366 | 0.480 |
| Peak Angle | **sex** | 2.059 | 0.357 | -2.515 | 6.633 |
| | **leg length** | -0.055 | 0.723 | -0.391 | 0.281 |
| | **race** | | | | |
| | Black | Ref. | Ref. | Ref. | |
| | Asian | 4.194 | 0.122 | -1.496 | 9.885 |
| | White | -1.375 | 0.588 | -7.03 | 4.280 |
| | Hispanic | 2.593 | 0.219 | -1.891 | 7.076 |
| | Mixed | 2.011 | 0.416 | -3.386 | 7.407 |
| | **speed** | 5.356 | 0.019* | 0.957 | 9.756 |
| *Conditional $R^2$* | 0.952 | | | | |
| *Marginal $R^2$* | 0.342 | | | | |
| Knee Excursion | (Intercept) | 21.90 | 0.086 | -3.714 | 47.52 |
| | **group** | -2.252 | 0.079 | -4.788 | 0.282 |
| | **age** | -0.068 | 0.713 | -0.456 | 0.320 |
| | **sex** | -0.984 | 0.612 | -4.987 | 3.018 |
| | **leg length** | -0.116 | 0.314 | -0.359 | 0.128 |
| | **race** | | | | |
| | Black | Ref. | Ref. | Ref. | |
| | Asian | 1.710 | 0.358 | -2.468 | 5.888 |
| | White | -0.451 | 0.829 | -5.142 | 4.240 |
| | Hispanic | 1.458 | 0.540 | -3.794 | 6.710 |
| | Mixed | 3.788 | 0.083 | -0.613 | 8.188 |
| | **speed** | 1.097 | 0.648 | -3.765 | 5.958 |
| *Conditional $R^2$* | 0.883 | | | | |

| | | | | | |
|---|---|---|---|---|---|
| *Marginal $R^2$* | 0.181 | | | | |
| 1st Peak Moment | (Intercept) | 0.772 | 0.279 | -0.725 | 2.268 |
| | **group** | -0.172 | **0.049*** | -0.344 | -0.001 |
| | **age** | -0.006 | 0.477 | -0.024 | 0.012 |
| | **sex** | 0.100 | 0.284 | -0.090 | 0.290 |
| | **leg length** | -0.004 | 0.527 | -0.020 | 0.011 |
| | **race** | | | | |
| | Black | Ref. | Ref. | Ref. | |
| | Asian | 0.135 | 0.184 | -0.084 | 0.355 |
| | White | -0.014 | 0.888 | -0.245 | 0.216 |
| | Hispanic | 0.243 | 0.019 | 0.051 | 0.434 |
| | Mixed | 0.216 | 0.118 | -0.068 | 0.499 |
| | **speed** | 0.322 | **0.024*** | 0.046 | 0.598 |
| *Conditional $R^2$* | 0.924 | | | | |
| *Marginal $R^2$* | 0.359 | | | | |
| 2nd Peak Moment | (Intercept) | -0.651 | 0.316 | -2.020 | 0.718 |
| | **group** | 0.245 | **0.003**** | 0.092 | 0.398 |
| | **age** | 0.01 | 0.053 | 0 | 0.020 |
| | **sex** | 0.02 | 0.754 | -0.114 | 0.155 |
| | **leg length** | 0 | 0.999 | -0.013 | 0.013 |
| | **race** | | | | |
| | Black | Ref. | Ref. | Ref. | |
| | Asian | 0.105 | 0.152 | -0.051 | 0.261 |
| | White | -0.009 | 0.900 | -0.175 | 0.157 |
| | Hispanic | 0.194 | 0.234 | -0.154 | 0.543 |
| | Mixed | 0.124 | 0.491 | -0.270 | 0.518 |
| | **speed** | 0.285 | 0.142 | -0.101 | 0.671 |
| *Conditional $R^2$* | 0.849 | | | | |
| *Marginal $R^2$* | 0.235 | | | | |
| Moment Range | (Intercept) | 0.400 | 0.613 | -1.301 | 2.100 |
| | **group** | -0.169 | **0.037*** | -0.326 | -0.011 |
| | **age** | -0.004 | 0.639 | -0.022 | 0.014 |
| | **sex** | 0.080 | 0.371 | -0.103 | 0.262 |
| | **leg length** | -0.002 | 0.838 | -0.019 | 0.016 |
| | **race** | | | | |
| | Black | Ref. | Ref. | Ref. | |
| | Asian | 0.147 | 0.152 | -0.071 | 0.364 |
| | White | 0.004 | 0.969 | -0.224 | 0.232 |
| | Hispanic | 0.246 | 0.075 | -0.031 | 0.523 |
| | Mixed | 0.204 | 0.101 | -0.050 | 0.457 |
| | **speed** | 0.527 | **<.001***** | 0.274 | 0.780 |
| *Conditional $R^2$* | 0.914 | | | | |
| *Marginal $R^2$* | 0.388 | | | | |
| **OCHB-TL** | (Intercept) | 18.045 | 0.347 | -22.55 | 58.64 |
| | **group** | -5.476 | **0.003**** | -8.864 | -2.088 |
| | **age** | 0.028 | 0.893 | -0.404 | 0.459 |
| Peak Angle | **sex** | 2.834 | 0.208 | -1.723 | 7.390 |
| | **leg length** | -0.069 | 0.674 | -0.423 | 0.286 |
| | **race** | | | | |
| | Black | Ref. | Ref. | Ref. | |
| | Asian | 4.673 | 0.109 | -1.38 | 10.73 |
| | White | -0.086 | 0.974 | -6.053 | 5.882 |
| | Hispanic | 2.576 | 0.300 | -2.782 | 7.933 |
| | Mixed | 2.786 | 0.280 | -2.748 | 8.320 |
| | **speed** | 1.916 | 0.270 | -1.578 | 5.410 |
| *Conditional $R^2$* | 0.946 | | | | |
| *Marginal $R^2$* | 0.340 | | | | |

| | | | | | |
|---|---|---|---|---|---|
| Knee Excursion | (Intercept) | 21.03 | 0.111 | -5.766 | 47.82 |
| | **group** | -3.452 | **0.006**** | -5.820 | -1.085 |
| | **age** | -0.067 | 0.711 | -0.443 | 0.310 |
| | **sex** | -0.423 | 0.824 | -4.365 | 3.520 |
| | **leg length** | -0.053 | 0.662 | -0.318 | 0.211 |
| | **race** | | | | |
| | Black | Ref. | Ref. | Ref. | |
| | Asian | 2.073 | 0.286 | -2.229 | 6.374 |
| | White | 0.721 | 0.735 | -4.042 | 5.484 |
| | Hispanic | 1.964 | 0.354 | -2.642 | 6.570 |
| | Mixed | 4.055 | 0.072 | -0.451 | 8.562 |
| | **speed** | -2.657 | 0.092 | -5.783 | 0.468 |
| *Conditional* $R^2$ | 0.865 | | | | |
| *Marginal* $R^2$ | 0.179 | | | | |
| 1st Peak Moment | (Intercept) | 0.968 | 0.207 | -0.625 | 2.561 |
| | **group** | -0.167 | **0.039*** | -0.325 | -0.009 |
| | **age** | -0.006 | 0.44 | -0.024 | 0.011 |
| | **sex** | 0.071 | 0.431 | -0.115 | 0.258 |
| | **leg length** | -0.007 | 0.37 | -0.022 | 0.009 |
| | **race** | | | | |
| | Black | Ref. | Ref. | Ref. | |
| | Asian | 0.130 | 0.231 | -0.107 | 0.366 |
| | White | 0.016 | 0.883 | -0.226 | 0.257 |
| | Hispanic | 0.218 | 0.059 | -0.011 | 0.446 |
| | Mixed | 0.176 | 0.163 | -0.088 | 0.440 |
| | **speed** | 0.363 | **<.001***** | 0.177 | 0.549 |
| *Conditional* $R^2$ | 0.912 | | | | |
| *Marginal* $R^2$ | 0.339 | | | | |
| 2nd Peak Moment | (Intercept) | 0.057 | 0.952 | -2.014 | 2.129 |
| | **group** | 0.211 | **0.039*** | 0.012 | 0.410 |
| | **age** | 0.013 | 0.077 | -0.002 | 0.027 |
| | **sex** | 0.054 | 0.525 | -0.121 | 0.229 |
| | **leg length** | -0.006 | 0.582 | -0.030 | 0.018 |
| | **race** | | | | |
| | Black | Ref. | Ref. | Ref. | |
| | Asian | 0.055 | 0.686 | -0.257 | 0.366 |
| | White | -0.059 | 0.659 | -0.360 | 0.242 |
| | Hispanic | 0.179 | 0.391 | -0.276 | 0.634 |
| | Mixed | 0.187 | 0.536 | -0.478 | 0.852 |
| | **speed** | 0.143 | 0.365 | -0.176 | 0.462 |
| *Conditional* $R^2$ | 0.908 | | | | |
| *Marginal* $R^2$ | 0.166 | | | | |
| Moment Range | (Intercept) | 0.744 | 0.372 | -1.024 | 2.511 |
| | **group** | -0.172 | **0.034*** | -0.33 | -0.014 |
| | **age** | -0.004 | 0.651 | -0.021 | 0.014 |
| | **sex** | 0.036 | 0.689 | -0.151 | 0.224 |
| | **leg length** | -0.005 | 0.526 | -0.023 | 0.013 |
| | **race** | | | | |
| | Black | Ref. | Ref. | Ref. | |
| | Asian | 0.151 | 0.185 | -0.094 | 0.395 |
| | White | 0.025 | 0.820 | -0.226 | 0.277 |
| | Hispanic | 0.225 | 0.113 | -0.066 | 0.516 |
| | Mixed | 0.175 | 0.157 | -0.083 | 0.434 |
| | **speed** | 0.571 | **<.001***** | 0.381 | 0.761 |
| *Conditional* $R^2$ | 0.906 | | | | |
| *Marginal* $R^2$ | 0.378 | | | | |
| **OCMA-LL** | (Intercept) | 32.04 | 0.290 | -31.73 | 95.82 |

| | | | | | |
|---|---|---|---|---|---|
| Peak Angle | **group** | -6.123 | **0.007**** | -10.40 | -1.851 |
| | **age** | 0.099 | 0.683 | -0.408 | 0.606 |
| | **sex** | 0.552 | 0.839 | -5.068 | 6.172 |
| | **leg length** | -0.432 | 0.183 | -1.104 | 0.239 |
| | **race** | | | | |
| | Black | Ref. | Ref. | Ref. | |
| | Asian | 3.892 | 0.349 | -5.405 | 13.19 |
| | White | 1.728 | 0.692 | -7.986 | 11.44 |
| | Hispanic | 3.927 | 0.443 | -7.256 | 15.11 |
| | Mixed | 1.753 | 0.667 | -7.222 | 10.73 |
| | **speed** | 13.61 | **0.007**** | 4.162 | 23.06 |
| *Conditional $R^2$* | 0.892 | | | | |
| *Marginal $R^2$* | 0.280 | | | | |
| Knee Excursion | (Intercept) | 36.29 | 0.052 | -0.460 | 73.04 |
| | **group** | -3.671 | **0.017*** | -6.620 | -0.723 |
| | **age** | -0.033 | 0.862 | -0.434 | 0.367 |
| | **sex** | -1.573 | 0.474 | -6.082 | 2.936 |
| | **leg length** | -0.408 | 0.051 | -0.818 | 0.002 |
| | **race** | | | | |
| | Black | Ref. | Ref. | Ref. | |
| | Asian | -0.476 | 0.862 | -6.805 | 5.854 |
| | White | 1.030 | 0.747 | -6.082 | 8.142 |
| | Hispanic | 0.546 | 0.879 | -7.454 | 8.547 |
| | Mixed | 4.100 | 0.220 | -2.956 | 11.16 |
| | **speed** | 5.886 | 0.084 | -0.847 | 12.62 |
| *Conditional $R^2$* | 0.836 | | | | |
| *Marginal $R^2$* | 0.251 | | | | |
| 1st Peak Moment | (Intercept) | 1.660 | 0.082 | -0.255 | 3.575 |
| | **group** | -0.234 | **0.010**** | -0.408 | -0.060 |
| | **age** | -0.005 | 0.608 | -0.024 | 0.015 |
| | **sex** | -0.022 | 0.827 | -0.234 | 0.190 |
| | **leg length** | -0.019 | 0.075 | -0.041 | 0.002 |
| | **race** | | | | |
| | Black | Ref. | Ref. | Ref. | |
| | Asian | 0.075 | 0.600 | -0.253 | 0.402 |
| | White | 0.056 | 0.728 | -0.302 | 0.413 |
| | Hispanic | 0.143 | 0.501 | -0.322 | 0.607 |
| | Mixed | 0.057 | 0.757 | -0.350 | 0.464 |
| | **speed** | 0.583 | **0.009*** | 0.159 | 1.008 |
| *Conditional $R^2$* | 0.882 | | | | |
| *Marginal $R^2$* | 0.286 | | | | |
| 2nd Peak Moment | (Intercept) | -0.846 | 0.054 | -1.709 | 0.017 |
| | **group** | 0.23 | **<.001***** | 0.113 | 0.346 |
| | **age** | 0.013 | 0.016 | 0.003 | 0.024 |
| | **sex** | 0.023 | 0.686 | -0.093 | 0.139 |
| | **leg length** | 0 | 0.951 | -0.010 | 0.010 |
| | **race** | | | | |
| | Black | Ref. | Ref. | Ref. | |
| | Asian | 0.099 | 0.240 | -0.035 | 0.233 |
| | White | 0.007 | 0.933 | -0.133 | 0.148 |
| | Hispanic | 0.126 | 0.232 | -0.035 | 0.288 |
| | Mixed | 0.052 | 0.630 | -0.131 | 0.234 |
| | **speed** | 0.441 | **<.001***** | 0.190 | 0.693 |
| *Conditional $R^2$* | 0.869 | | | | |
| *Marginal $R^2$* | 0.346 | | | | |
| | (Intercept) | 1.298 | 0.101 | -0.304 | 2.900 |
| | **group** | -0.198 | **0.012*** | -0.348 | -0.049 |

| | | | | | |
|---|---|---|---|---|---|
| Moment Range | **age** | -0.006 | 0.414 | -0.020 | 0.009 |
| | **sex** | -0.087 | 0.290 | -0.255 | 0.081 |
| | **leg length** | -0.016 | 0.081 | -0.033 | 0.002 |
| | **race** | | | | |
| | Black | Ref. | Ref. | Ref. | |
| | Asian | 0.024 | 0.811 | -0.208 | 0.256 |
| | White | -0.021 | 0.850 | -0.271 | 0.229 |
| | Hispanic | 0.010 | 0.949 | -0.320 | 0.339 |
| | Mixed | 0.038 | 0.799 | -0.294 | 0.370 |
|  | **speed** | 0.826 | **<.001***** | 0.386 | 1.266 |
| *Conditional $R^2$* | 0.828 | | | | |
| *Marginal $R^2$* | 0.370 | | | | |
| **OCHA-LL** | (Intercept) | 31.31 | 0.266 | -27.79 | 90.41 |
| | **group** | -7.895 | **0.002**** | -12.63 | -3.162 |
| | **age** | 0.096 | 0.735 | -0.495 | 0.687 |
| Peak Angle | **sex** | 0.282 | 0.927 | -6.093 | 6.657 |
| | **leg length** | -0.375 | 0.196 | -0.979 | 0.229 |
| | **race** | | | | |
| | Black | Ref. | Ref. | Ref. | |
| | Asian | 4.479 | 0.340 | -6.008 | 14.966 |
| | White | 3.285 | 0.514 | -7.846 | 14.416 |
| | Hispanic | 5.818 | 0.280 | -5.758 | 17.393 |
| | Mixed | 2.934 | 0.521 | -7.087 | 12.954 |
| | **speed** | 10.74 | **0.002**** | 4.201 | 17.285 |
| *Conditional $R^2$* | 0.888 | | | | |
| *Marginal $R^2$* | 0.294 | | | | |
| | (Intercept) | 32.90 | 0.078 | -4.378 | 70.19 |
| | **group** | -4.232 | **0.021*** | -7.776 | -0.688 |
| | **age** | -0.096 | 0.684 | -0.587 | 0.395 |
| Knee Excursion | **sex** | -1.227 | 0.651 | -6.820 | 4.366 |
| | **leg length** | -0.329 | 0.114 | -0.752 | 0.094 |
| | **race** | | | | |
| | Black | Ref. | Ref. | Ref. | |
| | Asian | -0.885 | 0.784 | -8.375 | 6.605 |
| | White | 1.760 | 0.658 | -7.115 | 10.64 |
| | Hispanic | 0.934 | 0.803 | -7.382 | 9.249 |
| | Mixed | 3.544 | 0.343 | -4.510 | 11.60 |
| | **speed** | 3.613 | 0.251 | -2.730 | 9.957 |
| *Conditional $R^2$* | 0.841 | | | | |
| *Marginal $R^2$* | 0.196 | | | | |
| | (Intercept) | 1.37 | 0.119 | -0.417 | 3.157 |
| | **group** | -0.28 | **0.006**** | -0.470 | -0.090 |
| | **age** | -0.003 | 0.773 | -0.025 | 0.019 |
| 1st Peak Moment | **sex** | 0.041 | 0.743 | -0.219 | 0.301 |
| | **leg length** | -0.016 | 0.091 | -0.034 | 0.003 |
| | **race** | | | | |
| | Black | Ref. | Ref. | Ref. | |
| | Asian | 0.119 | 0.429 | -0.222 | 0.461 |
| | White | 0.130 | 0.449 | -0.248 | 0.508 |
| | Hispanic | 0.177 | 0.353 | -0.237 | 0.592 |
| | Mixed | 0.157 | 0.361 | -0.214 | 0.528 |
| | **speed** | 0.434 | **0.014*** | 0.096 | 0.773 |
| *Conditional $R^2$* | 0.864 | | | | |
| *Marginal $R^2$* | 0.287 | | | | |
| | (Intercept) | -1.002 | **0.015*** | -1.764 | -0.240 |
| | **group** | 0.222 | **0.002**** | 0.088 | 0.357 |
| | **age** | 0.013 | **0.008**** | 0.004 | 0.023 |

| | | | | | |
|---|---|---|---|---|---|
| 2nd Peak Moment | **sex** | 0.012 | 0.831 | -0.107 | 0.132 |
| | **leg length** | 0.003 | 0.412 | -0.005 | 0.012 |
| | **race** | | | | |
| | Black | Ref. | Ref. | Ref. | |
| | Asian | 0.077 | 0.250 | -0.070 | 0.225 |
| | White | 0.008 | 0.915 | -0.159 | 0.175 |
| | Hispanic | 0.153 | 0.091 | -0.030 | 0.337 |
| | Mixed | 0.045 | 0.556 | -0.123 | 0.214 |
| | **speed** | 0.341 | **0.002**** | 0.142 | 0.540 |
| *Conditional $R^2$* | 0.890 | | | | |
| *Marginal $R^2$* | 0.350 | | | | |
| Moment Range | (Intercept) | 0.882 | 0.198 | -0.539 | 2.303 |
| | **group** | -0.272 | **0.003**** | -0.441 | -0.103 |
| | **age** | 0 | 0.983 | -0.017 | 0.017 |
| | **sex** | -0.021 | 0.845 | -0.245 | 0.203 |
| | **leg length** | -0.009 | 0.202 | -0.024 | 0.006 |
| | **race** | | | | |
| | Black | Ref. | Ref. | Ref. | |
| | Asian | 0.089 | 0.425 | -0.162 | 0.339 |
| | White | 0.067 | 0.580 | -0.201 | 0.335 |
| | Hispanic | 0.072 | 0.573 | -0.210 | 0.355 |
| | Mixed | 0.191 | 0.225 | -0.141 | 0.523 |
| | **speed** | 0.508 | **0.009**** | 0.138 | 0.879 |
| *Conditional $R^2$* | 0.808 | | | | |
| *Marginal $R^2$* | 0.326 | | | | |

Note: group– normal BMI group and obese group; PRF– Preferred speed walking; FF– Fast speed walking; OCMB-TL– Stance of trailing limb before Obstacle crossing medium height; OCHB-TL– Stance of trailing limb before Obstacle crossing high height; OCMA-LL– Stance of leading limb after Obstacle crossing medium height; OCHA-LL– Stance of leading limb after Obstacle crossing high height; Peak Angle– peak flexion angle during early stance; Knee Excursion– knee excursion from initial to peak (range of motion between heel-strike and midstance); 1st Peak Moment– peak knee extensor moment during early stance; 2nd Peak Moment– peak knee extensor moment during late stance; Moment Range – knee moment range from 1st flexion moment peak to1st extension moment peak; *β*– Coefficient; *CI*– Confidence interval; Conditional $R^2$– the variance explained by both fixed and random effects in the model; Marginal $R^2$– the variance explained by the fixed effects alone; *– Significance with $p < 0.05$; **– Significance with $p < 0.01$; ***– Significance with $p < 0.001$.

**Supplemental Table S4.** The effects of predictors on knee discrete outcomes in each task (PRF, FF, OCMB-TL, OCHB-TL, OCMA-LL and OCHA-LL) in female participants.

| | Source | $\beta$ | *p* | *95%CI lower for* $\beta$ | *95%CI upper for* $\beta$ |
|---|---|---|---|---|---|
| **PRF** | (Intercept) | 14.294 | 0.405 | -23.11 | 51.70 |
| | **fNB** | Ref. | Ref. | Ref. | |
| | **Apple** | -2.442 | 0.363 | -8.076 | 3.191 |
| | **Pear** | -6.83 | **0.004**** | -11.20 | -2.463 |
| Peak Angle | **age** | -0.046 | 0.877 | -0.688 | 0.595 |
| | **leg length** | -0.161 | 0.353 | -0.538 | 0.215 |
| | **race** | | | | |
| | Black | Ref. | Ref. | Ref. | |
| | Asian | 0.694 | 0.857 | -9.077 | 10.47 |
| | White | -4.383 | 0.284 | -14.32 | 5.548 |
| | Hispanic | 0.442 | 0.912 | -9.409 | 10.29 |
| | Mixed | -0.877 | 0.799 | -9.391 | 7.638 |
| | **speed** | 16.13 | **<.001***** | 11.61 | 20.65 |
| *Conditional* $R^2$ | 0.956 | | | | |
| *Marginal* $R^2$ | 0.330 | | | | |
| | (Intercept) | 19.13 | 0.150 | -8.542 | 46.79 |
| | **fNB** | Ref. | Ref. | Ref. | |
| | **Apple** | -2.119 | 0.336 | -6.725 | 2.487 |
| Knee Excursion | **Pear** | -2.666 | **0.150*** | -6.408 | 1.075 |
| | **age** | 0.094 | 0.691 | -0.414 | 0.603 |
| | **leg length** | -0.240 | 0.107 | -0.543 | 0.064 |
| | **race** | | | | |
| | Black | Ref. | Ref. | Ref. | |
| | Asian | -1.553 | 0.567 | -8.289 | 5.182 |
| | White | -4.070 | 0.094 | -9.231 | 1.090 |
| | Hispanic | -0.250 | 0.92 | -6.358 | 5.858 |
| | Mixed | 0.040 | 0.985 | -5.106 | 5.187 |
| | **speed** | 10.29 | **0.006**** | 3.367 | 17.21 |
| *Conditional* $R^2$ | 0.899 | | | | |
| *Marginal* $R^2$ | 0.207 | | | | |
| | (Intercept) | 0.846 | 0.293 | -0.883 | 2.574 |
| | **fNB** | Ref. | Ref. | Ref. | |
| | **Apple** | -0.072 | 0.477 | -0.287 | 0.142 |
| 1st Peak Moment | **Pear** | -0.110 | 0.204 | -0.287 | 0.067 |
| | **age** | 0 | 0.964 | -0.024 | 0.023 |
| | **leg length** | -0.016 | 0.063 | -0.033 | 0.001 |
| | **race** | | | | |
| | Black | Ref. | Ref. | Ref. | |
| | Asian | -0.006 | 0.971 | -0.447 | 0.434 |
| | White | -0.159 | 0.297 | -0.528 | 0.210 |
| | Hispanic | 0.143 | 0.404 | -0.261 | 0.547 |
| | Mixed | 0.207 | 0.259 | -0.212 | 0.625 |
| | **speed** | 0.920 | **<.001***** | 0.680 | 1.161 |
| *Conditional* $R^2$ | 0.919 | | | | |
| *Marginal* $R^2$ | 0.382 | | | | |
| | (Intercept) | -0.816 | 0.056 | -1.658 | 0.026 |
| | **fNB** | Ref. | Ref. | Ref. | |
| | **Apple** | 0.197 | 0.056 | -0.006 | 0.400 |
| 2nd Peak Moment | **Pear** | 0.135 | 0.075 | -0.015 | 0.286 |
| | **age** | 0.007 | 0.296 | -0.007 | 0.022 |

| | | | | | |
|---|---|---|---|---|---|
| | **leg length** | 0.003 | 0.548 | -0.007 | 0.012 |
| | **race** | | | | |
| | Black | Ref. | Ref. | Ref. | |
| | Asian | 0.091 | 0.201 | -0.071 | 0.253 |
| | White | 0.067 | 0.304 | -0.092 | 0.226 |
| | Hispanic | 0.114 | 0.130 | -0.049 | 0.277 |
| | Mixed | 0 | 1 | -0.183 | 0.183 |
| | **speed** | 0.367 | **0.001**** | 0.175 | 0.559 |
| *Conditional $R^2$* | 0.927 | | | | |
| *Marginal $R^2$* | 0.320 | | | | |
| | (Intercept) | 0.500 | 0.497 | -1.115 | 2.115 |
| | **fNB** | Ref. | Ref. | Ref. | |
| | **Apple** | -0.085 | 0.338 | -0.271 | 0.101 |
| | **Pear** | -0.121 | 0.135 | -0.285 | 0.042 |
| Moment Range | **age** | 0 | 0.992 | -0.022 | 0.022 |
| | **leg length** | -0.014 | 0.077 | -0.030 | 0.002 |
| | **race** | | | | |
| | Black | Ref. | Ref. | Ref. | |
| | Asian | -0.104 | 0.512 | -0.495 | 0.286 |
| | White | -0.214 | 0.144 | -0.541 | 0.113 |
| | Hispanic | 0.050 | 0.736 | -0.310 | 0.410 |
| | Mixed | 0.068 | 0.642 | -0.285 | 0.420 |
| | **speed** | 1.257 | **<.001***** | 0.998 | 1.516 |
| *Conditional $R^2$* | 0.929 | | | | |
| *Marginal $R^2$* | 0.267 | | | | |
| **FF** | (Intercept) | 16.61 | 0.407 | -27.04 | 60.26 |
| | **fNB** | Ref. | Ref. | Ref. | |
| | **Apple** | -2.441 | 0.485 | -9.835 | 4.953 |
| Peak Angle | **Pear** | -8.870 | **0.004**** | -14.546 | -3.195 |
| | **age** | -0.003 | 0.993 | -0.664 | 0.659 |
| | **leg length** | 0.030 | 0.872 | -0.388 | 0.448 |
| | **race** | | | | |
| | Black | Ref. | Ref. | Ref. | |
| | Asian | 3.192 | 0.504 | -8.729 | 15.11 |
| | White | -1.869 | 0.701 | -14.69 | 10.96 |
| | Hispanic | 2.320 | 0.616 | -8.986 | 13.63 |
| | Mixed | 4.589 | 0.327 | -6.402 | 15.58 |
| | **speed** | 2.668 | 0.091 | -0.478 | 5.814 |
| *Conditional $R^2$* | 0.943 | | | | |
| *Marginal $R^2$* | 0.286 | | | | |
| | (Intercept) | 24.21 | 0.077 | -3.253 | 51.67 |
| | **fNB** | Ref. | Ref. | Ref. | |
| | **Apple** | -2.453 | 0.278 | -7.158 | 2.252 |
| Knee Excursion | **Pear** | -4.006 | **0.033*** | -7.638 | -0.373 |
| | **age** | 0.233 | 0.235 | -0.175 | 0.641 |
| | **leg length** | -0.167 | 0.152 | -0.410 | 0.077 |
| | **race** | | | | |
| | Black | Ref. | Ref. | Ref. | |
| | Asian | 0.456 | 0.835 | -5.162 | 6.075 |
| | White | -1.865 | 0.361 | -6.959 | 3.229 |
| | Hispanic | 1.349 | 0.553 | -4.180 | 6.878 |
| | Mixed | 4.113 | 0.092 | -0.991 | 9.217 |
| | **speed** | -0.060 | 0.966 | -3.039 | 2.918 |
| *Conditional $R^2$* | 0.872 | | | | |
| *Marginal $R^2$* | 0.225 | | | | |
| | (Intercept) | 0.530 | 0.460 | -1.039 | 2.099 |
| | **fNB** | Ref. | Ref. | Ref. | |
| | **Apple** | -0.103 | 0.476 | -0.408 | 0.202 |

| | | | | | |
|---|---|---|---|---|---|
| 1st Peak Moment | **Pear** | -0.285 | **0.028*** | -0.536 | -0.035 |
| | **age** | 0.005 | 0.637 | -0.016 | 0.026 |
| | **leg length** | -0.004 | 0.552 | -0.021 | 0.012 |
| | **race** | | | | |
| | Black | Ref. | Ref. | Ref. | |
| | Asian | 0.245 | 0.114 | -0.090 | 0.580 |
| | White | 0.028 | 0.786 | -0.248 | 0.305 |
| | Hispanic | 0.328 | **0.042*** | 0.019 | 0.637 |
| | Mixed | 0.403 | 0.068 | -0.043 | 0.850 |
| | **speed** | 0.388 | **0.001**** | 0.196 | 0.580 |
| *Conditional $R^2$* | 0.896 | | | | |
| *Marginal $R^2$* | 0.393 | | | | |
| 2nd Peak Moment | (Intercept) | -1.507 | 0.011 | -2.557 | -0.456 |
| | **fNB** | Ref. | Ref. | Ref. | |
| | **Apple** | 0.242 | **0.035*** | 0.020 | 0.463 |
| | **Pear** | 0.197 | **0.028*** | 0.025 | 0.370 |
| | **age** | 0.014 | 0.091 | -0.003 | 0.030 |
| | **leg length** | 0.008 | 0.113 | -0.002 | 0.019 |
| | **race** | | | | |
| | Black | Ref. | Ref. | Ref. | |
| | Asian | 0.021 | 0.770 | -0.163 | 0.205 |
| | White | -0.054 | 0.476 | -0.248 | 0.140 |
| | Hispanic | 0.106 | 0.318 | -0.142 | 0.353 |
| | Mixed | -0.059 | 0.450 | -0.248 | 0.129 |
| | **speed** | 0.383 | **0.001**** | 0.193 | 0.574 |
| *Conditional $R^2$* | 0.882 | | | | |
| *Marginal $R^2$* | 0.328 | | | | |
| Moment Range | (Intercept) | 0.054 | 0.926 | -1.224 | 1.331 |
| | **fNB** | Ref. | Ref. | Ref. | |
| | **Apple** | -0.168 | 0.132 | -0.395 | 0.059 |
| | **Pear** | -0.351 | **0.002**** | -0.547 | -0.154 |
| | **age** | 0.007 | 0.437 | -0.012 | 0.026 |
| | **leg length** | -0.002 | 0.794 | -0.017 | 0.013 |
| | **race** | | | | |
| | Black | Ref. | Ref. | Ref. | |
| | Asian | 0.157 | 0.129 | -0.070 | 0.384 |
| | White | -0.088 | 0.190 | -0.244 | 0.068 |
| | Hispanic | 0.241 | **0.030** | 0.034 | 0.448 |
| | Mixed | 0.253 | **0.016** | 0.070 | 0.436 |
| | **speed** | 0.739 | **<.001***** | 0.532 | 0.947 |
| *Conditional $R^2$* | 0.860 | | | | |
| *Marginal $R^2$* | 0.519 | | | | |
| **OCMB-TL** Peak Angle | (Intercept) | 16.06 | 0.281 | -15.90 | 48.02 |
| | **fNB** | Ref. | Ref. | Ref. | |
| | **Apple** | -2.948 | 0.315 | -9.072 | 3.177 |
| | **Pear** | -8.326 | **0.002**** | -12.93 | -3.724 |
| | **age** | -0.013 | 0.961 | -0.578 | 0.552 |
| | **leg length** | -0.048 | 0.734 | -0.363 | 0.267 |
| | **race** | | | | |
| | Black | Ref. | Ref. | Ref. | |
| | Asian | 2.503 | 0.363 | -4.161 | 9.168 |
| | White | -2.251 | 0.328 | -7.928 | 3.426 |
| | Hispanic | 1.794 | 0.516 | -4.861 | 8.449 |
| | Mixed | -0.114 | 0.967 | -6.823 | 6.595 |
| | **speed** | 6.284 | **0.024*** | 0.903 | 11.67 |
| *Conditional $R^2$* | 0.952 | | | | |
| *Marginal $R^2$* | 0.342 | | | | |

| | | | | | |
|---|---|---|---|---|---|
| Knee Excursion | (Intercept) | 22.27 | 0.114 | -6.687 | 51.22 |
| | **fNB** | Ref. | Ref. | Ref. | |
| | **Apple** | -3.065 | 0.220 | -8.226 | 2.095 |
| | **Pear** | -3.867 | **0.047*** | -7.674 | -0.060 |
| | **age** | 0.147 | 0.498 | -0.313 | 0.606 |
| | **leg length** | -0.204 | 0.103 | -0.460 | 0.052 |
| | **race** | | | | |
| | Black | Ref. | Ref. | Ref. | |
| | Asian | 0.184 | 0.947 | -6.841 | 7.209 |
| | White | -1.683 | 0.414 | -6.855 | 3.490 |
| | Hispanic | 0.478 | 0.864 | -6.383 | 7.339 |
| | Mixed | 3.057 | 0.242 | -2.881 | 8.994 |
| | **speed** | 3.196 | 0.284 | -2.820 | 9.212 |
| *Conditional $R^2$* | 0.883 | | | | |
| *Marginal $R^2$* | 0.181 | | | | |
| 1st Peak Moment | (Intercept) | 0.803 | 0.343 | -1.031 | 2.636 |
| | **fNB** | Ref. | Ref. | Ref. | |
| | **Apple** | -0.163 | 0.315 | -0.501 | 0.175 |
| | **Pear** | -0.291 | **0.044*** | -0.573 | -0.009 |
| | **age** | -0.002 | 0.863 | -0.031 | 0.026 |
| | **leg length** | -0.003 | 0.710 | -0.024 | 0.017 |
| | **race** | | | | |
| | Black | Ref. | Ref. | Ref. | |
| | Asian | 0.163 | 0.327 | -0.237 | 0.563 |
| | White | 0.035 | 0.772 | -0.283 | 0.354 |
| | Hispanic | 0.296 | 0.072 | -0.039 | 0.631 |
| | Mixed | 0.284 | 0.159 | -0.159 | 0.727 |
| | **speed** | 0.226 | 0.189 | -0.119 | 0.571 |
| *Conditional $R^2$* | 0.924 | | | | |
| *Marginal $R^2$* | 0.359 | | | | |
| 2nd Peak Moment | (Intercept) | -0.841 | 0.237 | -2.356 | 0.673 |
| | **fNB** | Ref. | Ref. | Ref. | |
| | **Apple** | 0.339 | **0.048*** | 0.003 | 0.674 |
| | **Pear** | 0.189 | 0.123 | -0.057 | 0.436 |
| | **age** | 0.005 | 0.608 | -0.016 | 0.026 |
| | **leg length** | 0.007 | 0.376 | -0.010 | 0.023 |
| | **race** | | | | |
| | Black | Ref. | Ref. | Ref. | |
| | Asian | 0.182 | 0.277 | -0.213 | 0.577 |
| | White | 0.037 | 0.801 | -0.348 | 0.422 |
| | Hispanic | 0.268 | 0.173 | -0.167 | 0.704 |
| | Mixed | 0.187 | 0.517 | -0.504 | 0.878 |
| | **speed** | 0.045 | 0.877 | -0.552 | 0.643 |
| *Conditional $R^2$* | 0.849 | | | | |
| *Marginal $R^2$* | 0.235 | | | | |
| Moment Range | (Intercept) | 0.341 | 0.714 | -1.726 | 2.408 |
| | **fNB** | Ref. | Ref. | Ref. | |
| | **Apple** | -0.161 | 0.306 | -0.489 | 0.167 |
| | **Pear** | -0.299 | **0.035*** | -0.572 | -0.025 |
| | **age** | -0.001 | 0.963 | -0.030 | 0.028 |
| | **leg length** | 0.001 | 0.942 | -0.023 | 0.024 |
| | **race** | | | | |
| | Black | Ref. | Ref. | Ref. | |
| | Asian | 0.119 | 0.461 | -0.278 | 0.516 |
| | White | 0.024 | 0.845 | -0.296 | 0.343 |
| | Hispanic | 0.257 | 0.145 | -0.127 | 0.642 |
| | Mixed | 0.213 | 0.225 | -0.185 | 0.611 |

| | | | | | |
|---|---|---|---|---|---|
| | **speed** | 0.437 | **0.015*** | 0.094 | 0.780 |
| *Conditional $R^2$* | 0.914 | | | | |
| *Marginal $R^2$* | 0.388 | | | | |
| **OCHB-TL** | (Intercept) | 26.16 | 0.148 | -11.48 | 63.80 |
| | **fNB** | Ref. | Ref. | Ref. | |
| | **Apple** | -3.355 | 0.290 | -9.955 | 3.245 |
| Peak Angle | **Pear** | -8.086 | **0.002**** | -12.81 | -3.368 |
| | **age** | -0.030 | 0.925 | -0.707 | 0.648 |
| | **leg length** | -0.124 | 0.441 | -0.476 | 0.229 |
| | **race** | | | | |
| | Black | Ref. | Ref. | Ref. | |
| | Asian | 4.260 | 0.268 | -4.831 | 13.35 |
| | White | 0.346 | 0.915 | -8.285 | 8.978 |
| | Hispanic | 2.904 | 0.466 | -6.669 | 12.48 |
| | Mixed | 1.710 | 0.602 | -6.288 | 9.707 |
| | **speed** | 2.925 | 0.126 | -0.894 | 6.744 |
| *Conditional $R^2$* | 0.946 | | | | |
| *Marginal $R^2$* | 0.340 | | | | |
| | (Intercept) | 23.48 | 0.093 | -4.888 | 51.84 |
| | **fNB** | Ref. | Ref. | Ref. | |
| | **Apple** | -4.187 | 0.119 | -9.616 | 1.242 |
| Knee Excursion | **Pear** | -4.526 | **0.022*** | -8.318 | -0.733 |
| | **age** | 0.132 | 0.558 | -0.348 | 0.612 |
| | **leg length** | -0.157 | 0.244 | -0.444 | 0.131 |
| | **race** | | | | |
| | Black | Ref. | Ref. | Ref. | |
| | Asian | 1.270 | 0.662 | -6.075 | 8.616 |
| | White | 0.399 | 0.856 | -5.430 | 6.228 |
| | Hispanic | 1.654 | 0.569 | -5.398 | 8.706 |
| | Mixed | 3.898 | 0.166 | -2.298 | 10.06 |
| | **speed** | -0.993 | 0.564 | -4.526 | 2.540 |
| *Conditional $R^2$* | 0.865 | | | | |
| *Marginal $R^2$* | 0.179 | | | | |
| | (Intercept) | 1.112 | 0.221 | -0.818 | 3.041 |
| | **fNB** | Ref. | Ref. | Ref. | |
| | **Apple** | -0.143 | 0.323 | -0.445 | 0.159 |
| 1st Peak Moment | **Pear** | -0.229 | 0.071 | -0.481 | 0.022 |
| | **age** | -0.005 | 0.733 | -0.033 | 0.024 |
| | **leg length** | -0.009 | 0.368 | -0.031 | 0.013 |
| | **race** | | | | |
| | Black | Ref. | Ref. | Ref. | |
| | Asian | 0.154 | 0.327 | -0.224 | 0.533 |
| | White | 0.090 | 0.466 | -0.224 | 0.403 |
| | Hispanic | 0.272 | 0.086 | -0.057 | 0.600 |
| | Mixed | 0.237 | 0.167 | -0.141 | 0.614 |
| | **speed** | 0.404 | **0.001**** | 0.191 | 0.617 |
| *Conditional $R^2$* | 0.912 | | | | |
| *Marginal $R^2$* | 0.339 | | | | |
| | (Intercept) | -0.040 | 0.965 | -2.103 | 2.023 |
| | **fNB** | Ref. | Ref. | Ref. | |
| | **Apple** | 0.415 | 0.072 | -0.044 | 0.874 |
| 2nd Peak Moment | **Pear** | 0.166 | 0.238 | -0.121 | 0.452 |
| | **age** | 0.005 | 0.668 | -0.021 | 0.031 |
| | **leg length** | -0.003 | 0.777 | -0.028 | 0.022 |
| | **race** | | | | |
| | Black | Ref. | Ref. | Ref. | |

| | | | | | |
|---|---|---|---|---|---|
| | Asian | 0.231 | 0.226 | -0.212 | 0.674 |
| | White | 0.047 | 0.767 | -0.376 | 0.471 |
| | Hispanic | 0.331 | 0.145 | -0.164 | 0.826 |
| | Mixed | 0.299 | 0.428 | -0.601 | 1.199 |
| | **speed** | 0.070 | 0.732 | -0.350 | 0.489 |
| *Conditional $R^2$* | 0.908 | | | | |
| *Marginal $R^2$* | 0.166 | | | | |
| | (Intercept) | 0.777 | 0.414 | -1.299 | 2.853 |
| | **fNB** | | | | |
| | **Apple** | -0.149 | 0.325 | -0.467 | 0.168 |
| Moment Range | **Pear** | -0.241 | 0.073 | -0.508 | 0.025 |
| | **age** | -0.004 | 0.782 | -0.034 | 0.026 |
| | **leg length** | -0.006 | 0.608 | -0.030 | 0.019 |
| | **race** | | | | |
| | Black | Ref. | Ref. | Ref. | |
| | Asian | 0.116 | 0.429 | -0.244 | 0.477 |
| | White | 0.076 | 0.517 | -0.225 | 0.376 |
| | Hispanic | 0.238 | 0.171 | -0.147 | 0.622 |
| | Mixed | 0.186 | 0.229 | -0.165 | 0.536 |
| | **speed** | 0.602 | **<.001*****| 0.385 | 0.819 |
| *Conditional $R^2$* | 0.906 | | | | |
| *Marginal $R^2$* | 0.378 | | | | |
| **OCMA-LL** | (Intercept) | 30.82 | 0.320 | -36.12 | 97.76 |
| | **fNB** | Ref. | Ref. | Ref. | |
| | **Apple** | -7.374 | 0.120 | -16.97 | 2.225 |
| Peak Angle | **Pear** | -9.950 | **0.006**** | -16.63 | -3.273 |
| | **age** | 0.247 | 0.558 | -0.651 | 1.145 |
| | **leg length** | -0.468 | 0.218 | -1.271 | 0.335 |
| | **race** | | | | |
| | Black | Ref. | Ref. | Ref. | |
| | Asian | 1.444 | 0.824 | -15.08 | 17.97 |
| | White | 0.103 | 0.988 | -17.54 | 17.74 |
| | Hispanic | 2.583 | 0.718 | -14.90 | 20.06 |
| | Mixed | 1.748 | 0.784 | -13.67 | 17.16 |
| | **speed** | 16.82 | **0.011*** | 4.435 | 29.20 |
| *Conditional $R^2$* | 0.892 | | | | |
| *Marginal $R^2$* | 0.280 | | | | |
| | (Intercept) | 34.64 | 0.100 | -8.314 | 77.59 |
| | **fNB** | Ref. | Ref. | Ref. | |
| | **Apple** | -4.293 | 0.142 | -10.24 | 1.655 |
| Knee Excursion | **Pear** | -5.074 | **0.019*** | -9.185 | -0.963 |
| | **age** | 0.094 | 0.721 | -0.471 | 0.660 |
| | **leg length** | -0.425 | 0.085 | -0.922 | 0.072 |
| | **race** | | | | |
| | Black | Ref. | Ref. | Ref. | |
| | Asian | 0.650 | 0.890 | -11.30 | 12.60 |
| | White | 1.291 | 0.784 | -10.92 | 13.50 |
| | Hispanic | 1.625 | 0.750 | -10.82 | 14.06 |
| | Mixed | 6.631 | 0.202 | -4.836 | 18.10 |
| | **speed** | 4.415 | 0.314 | -4.593 | 13.42 |
| *Conditional $R^2$* | 0.836 | | | | |
| *Marginal $R^2$* | 0.251 | | | | |
| | (Intercept) | 1.345 | 0.176 | -0.739 | 3.429 |
| | **fNB** | Ref. | Ref. | Ref. | |
| | **Apple** | -0.35 | **0.042*** | -0.685 | -0.015 |
| 1st Peak Moment | **Pear** | -0.371 | **0.009**** | -0.633 | -0.109 |

| | | | | | |
|---|---|---|---|---|---|
| | **age** | 0.006 | 0.663 | -0.025 | 0.038 |
| | **leg length** | -0.02 | 0.099 | -0.045 | 0.005 |
| | **race** | | | | |
| | Black | Ref. | Ref. | Ref. | |
| | Asian | 0.051 | 0.821 | -0.516 | 0.617 |
| | White | 0.067 | 0.766 | -0.516 | 0.649 |
| | Hispanic | 0.158 | 0.552 | -0.483 | 0.800 |
| | Mixed | 0.203 | 0.388 | -0.344 | 0.750 |
| | **speed** | 0.680 | **0.018*** | 0.133 | 1.227 |
| *Conditional $R^2$* | 0.882 | | | | |
| *Marginal $R^2$* | 0.286 | | | | |
| 2nd Peak Moment | (Intercept) | -0.926 | 0.032 | -1.752 | -0.101 |
| | **fNB** | Ref. | Ref. | Ref. | |
| | **Apple** | 0.351 | **0.004**** | 0.137 | 0.565 |
| | **Pear** | 0.241 | **0.003**** | 0.093 | 0.388 |
| | **age** | 0.010 | 0.172 | -0.005 | 0.026 |
| | **leg length** | 0 | 0.999 | -0.011 | 0.011 |
| | **race** | | | | |
| | Black | Ref. | Ref. | Ref. | |
| | Asian | 0.148 | 0.142 | -0.074 | 0.371 |
| | White | 0.017 | 0.813 | -0.169 | 0.202 |
| | Hispanic | 0.164 | 0.096 | -0.042 | 0.369 |
| | Mixed | 0.051 | 0.679 | -0.242 | 0.344 |
| | **speed** | 0.507 | **0.008**** | 0.151 | 0.864 |
| *Conditional $R^2$* | 0.869 | | | | |
| *Marginal $R^2$* | 0.346 | | | | |
| Moment Range | (Intercept) | 1.096 | 0.157 | -0.521 | 2.712 |
| | **fNB** | Ref. | Ref. | Ref. | |
| | **Apple** | -0.3 | 0.052 | -0.604 | 0.003 |
| | **Pear** | -0.326 | **0.004**** | -0.532 | -0.120 |
| | **age** | -0.001 | 0.949 | -0.025 | 0.024 |
| | **leg length** | -0.016 | 0.083 | -0.035 | 0.003 |
| | **race** | | | | |
| | Black | Ref. | Ref. | Ref. | |
| | Asian | -0.078 | 0.606 | -0.452 | 0.297 |
| | White | -0.067 | 0.672 | -0.475 | 0.340 |
| | Hispanic | -0.046 | 0.802 | -0.488 | 0.397 |
| | Mixed | 0.066 | 0.682 | -0.317 | 0.450 |
| | **speed** | 0.954 | **0.001***** | 0.445 | 1.462 |
| *Conditional $R^2$* | 0.828 | | | | |
| *Marginal $R^2$* | 0.370 | | | | |
| **OCHA-LL** Peak Angle | (Intercept) | 21.67 | 0.402 | -34.70 | 78.03 |
| | **fNB** | Ref. | Ref. | Ref. | |
| | **Apple** | -9.700 | **0.035*** | -18.62 | -0.783 |
| | **Pear** | -13.36 | **0.001**** | -20.29 | -6.433 |
| | **age** | 0.380 | 0.357 | -0.488 | 1.247 |
| | **leg length** | -0.317 | 0.289 | -0.961 | 0.326 |
| | **race** | | | | |
| | Black | Ref. | Ref. | Ref. | |
| | Asian | 2.043 | 0.756 | -14.66 | 18.74 |
| | White | 1.726 | 0.795 | -15.77 | 19.22 |
| | Hispanic | 4.750 | 0.51 | -12.62 | 22.11 |
| | Mixed | 3.611 | 0.580 | -12.12 | 19.34 |
| | **speed** | 11.78 | **0.004**** | 4.459 | 19.09 |
| *Conditional $R^2$* | 0.888 | | | | |
| *Marginal $R^2$* | 0.294 | | | | |

| | | | | | |
|---|---|---|---|---|---|
| Knee Excursion | (Intercept) | 28.63 | 0.151 | -12.85 | 70.12 |
| | **fNB** | Ref. | Ref. | Ref. | |
| | **Apple** | -5.919 | **0.049*** | -11.79 | -0.044 |
| | **Pear** | -6.349 | **0.020*** | -11.55 | -1.145 |
| | **age** | 0.171 | 0.521 | -0.396 | 0.739 |
| | **leg length** | -0.365 | 0.120 | -0.847 | 0.118 |
| | **race** | | | | |
| | Black | Ref. | Ref. | Ref. | |
| | Asian | -1.053 | 0.842 | -14.46 | 12.35 |
| | White | 1.147 | 0.823 | -12.34 | 14.63 |
| | Hispanic | 1.104 | 0.840 | -12.32 | 14.52 |
| | Mixed | 5.727 | 0.310 | -7.233 | 18.69 |
| | **speed** | 3.927 | 0.262 | -3.216 | 11.07 |
| *Conditional $R^2$* | 0.841 | | | | |
| *Marginal $R^2$* | 0.196 | | | | |
| 1st Peak Moment | (Intercept) | 1.051 | 0.155 | -0.490 | 2.591 |
| | **fNB** | Ref. | Ref. | Ref. | |
| | **Apple** | -0.438 | **0.007**** | -0.730 | -0.146 |
| | **Pear** | -0.468 | **0.001**** | -0.714 | -0.221 |
| | **age** | 0.014 | 0.307 | -0.015 | 0.044 |
| | **leg length** | -0.017 | 0.075 | -0.035 | 0.002 |
| | **race** | | | | |
| | Black | Ref. | Ref. | Ref. | |
| | Asian | 0.066 | 0.743 | -0.445 | 0.577 |
| | White | 0.100 | 0.613 | -0.412 | 0.612 |
| | Hispanic | 0.168 | 0.481 | -0.404 | 0.739 |
| | Mixed | 0.280 | 0.193 | -0.196 | 0.757 |
| | **speed** | 0.522 | **0.008**** | 0.157 | 0.887 |
| *Conditional $R^2$* | 0.864 | | | | |
| *Marginal $R^2$* | 0.287 | | | | |
| 2nd Peak Moment | (Intercept) | -1.244 | **0.007**** | -2.043 | -0.445 |
| | **fNB** | Ref. | Ref. | Ref. | |
| | **Apple** | 0.307 | **0.018*** | 0.062 | 0.552 |
| | **Pear** | 0.221 | **0.021*** | 0.039 | 0.403 |
| | **age** | 0.012 | 0.177 | -0.006 | 0.029 |
| | **leg length** | 0.006 | 0.181 | -0.004 | 0.016 |
| | **race** | | | | |
| | Black | Ref. | Ref. | Ref. | |
| | Asian | 0.116 | 0.222 | -0.104 | 0.336 |
| | White | 0.042 | 0.627 | -0.184 | 0.269 |
| | Hispanic | 0.197 | 0.091 | -0.046 | 0.439 |
| | Mixed | 0.082 | 0.447 | -0.174 | 0.338 |
| | **speed** | 0.320 | **0.007**** | 0.100 | 0.540 |
| *Conditional $R^2$* | 0.890 | | | | |
| *Marginal $R^2$* | 0.350 | | | | |
| Moment Range | (Intercept) | 0.644 | 0.204 | -0.428 | 1.717 |
| | **fNB** | Ref. | Ref. | Ref. | |
| | **Apple** | -0.414 | **0.002**** | -0.650 | -0.178 |
| | **Pear** | -0.458 | **<.001***** | -0.625 | -0.290 |
| | **age** | 0.011 | 0.245 | -0.009 | 0.031 |
| | **leg length** | -0.010 | 0.102 | -0.022 | 0.002 |
| | **race** | | | | |
| | Black | Ref. | Ref. | Ref. | |
| | Asian | -0.017 | 0.895 | -0.351 | 0.316 |
| | White | 0.005 | 0.966 | -0.329 | 0.340 |
| | Hispanic | 0.014 | 0.928 | -0.376 | 0.405 |
| | Mixed | 0.228 | 0.17 | -0.131 | 0.587 |

| | | | | | |
|---|---|---|---|---|---|
| | **speed** | 0.628 | 0.005 | 0.214 | 1.042 |
| *Conditional $R^2$* | 0.808 | | | | |
| *Marginal $R^2$* | 0.326 | | | | |

Note: fNB– female participants with normal BMI; Apple– Apple shape; Pear– Pear shape; PRF– Preferred speed walking; FF– Fast speed walking; OCMB-TL– Stance of trailing limb before Obstacle crossing medium height; OCHB-TL– Stance of trailing limb before Obstacle crossing high height; OCMA-LL – Stance of leading limb after Obstacle crossing medium height; OCHA-LL– Stance of leading limb after Obstacle crossing high height; Peak Angle– peak flexion angle during early stance; Knee Excursion– knee excursion from initial to peak (range of motion between heel-strike and midstance); 1st Peak Moment– peak knee extensor moment during early stance; 2nd Peak Moment– peak knee extensor moment during late stance; Moment Range – knee moment range from 1st flexion moment peak to1st extension moment peak; $\beta$– Coefficient; *CI*– Confidence interval; Conditional $R^2$ – the variance explained by both fixed and random effects in the model; Marginal $R^2$ – the variance explained by the fixed effects alone; * – Significance with $p<0.05$; ** – Significance with $p<0.01$; *** – Significance with $p<0.001$.

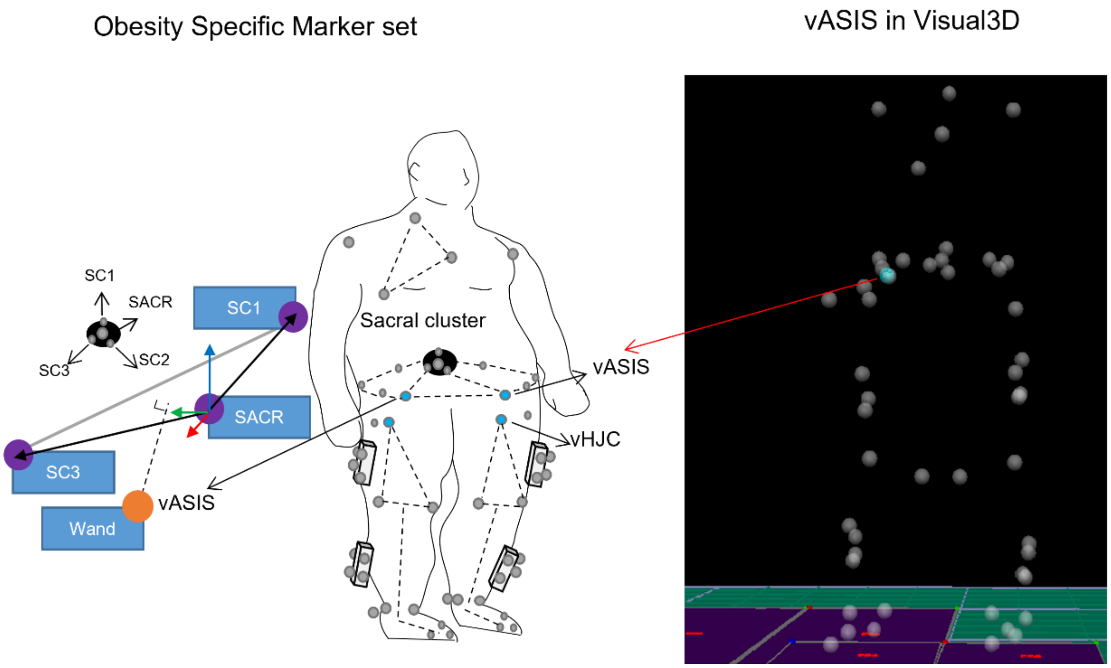


**Fig. S1** — Placement of reflective markers with sacral, thigh, and shank clusters and reconstructed virtual ASIS (vASIS) in Visual3D.

Note: ASIS– Anterior Superior Iliac Spine; SACR– Sacral maker; SC1– Sacral Cluster Superior; SC2– Sacral Cluster Inferior; SC3– Sacral Cluster Lateral; vHJC– virtual Hip Joint Center.

.

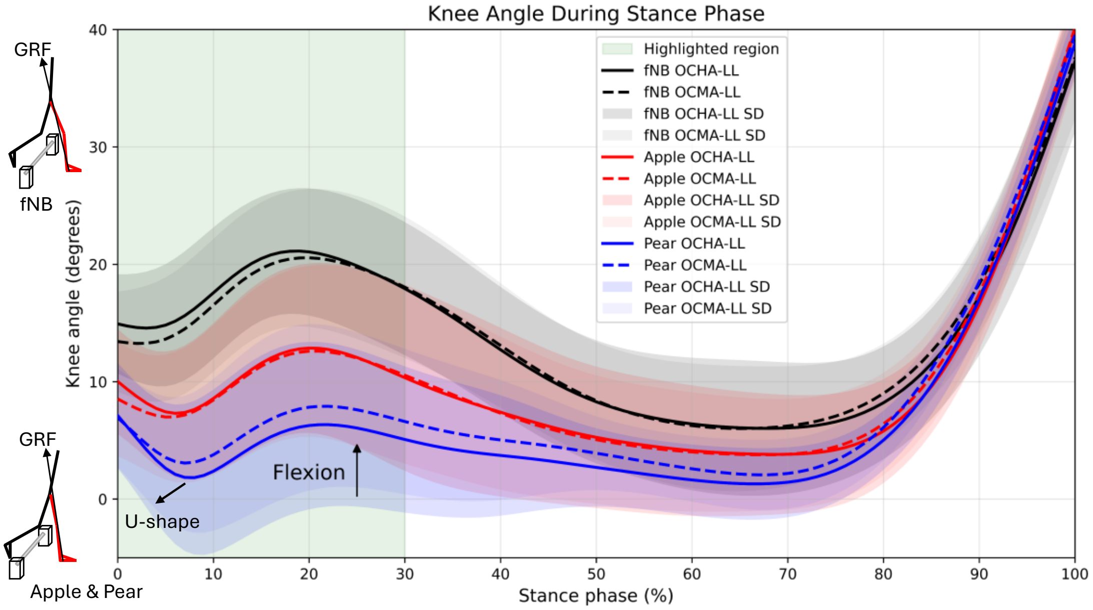


**Fig. S2** — Knee flexion angle during OCMA-LL and OCHA-LL in female participants with normal BMI (fNB, black), apple shape (Apple, red), and pear shape (Pear, blue). Apple and Pear showed a U-shaped early-stance waveform, in which knee flexion decreased shortly after heel strike and then returned toward the initial contact angle, rather than the larger flexion excursion seen in fNB during the highlighted weight-acceptance region (light green). This straighter-knee pattern may align the vertical and anterior–posterior ground reaction forces more closely with the knee joint center, decreasing the external knee-flexion moment. In contrast, the greater early-stance flexion in fNB places the ground reaction force further behind the knee, increasing the external knee-flexion moment. To counteract each flexion moment, the knee extensors must generate a corresponding internal extension moment of equal magnitude.

Note: OCMA-LL– Stance of leading limb after Obstacle crossing medium height; OCHA-LL– Stance of leading limb after Obstacle crossing high height; SD– standard deviation.
